# Synthetic Growth Hormone-Releasing Hormone Agonist as Novel Treatment for Heart Failure with Preserved Ejection Fraction

**DOI:** 10.1101/2020.02.28.967000

**Authors:** Raul A. Dulce, Rosemeire M. Kanashiro-Takeuchi, Lauro M. Takeuchi, Alessandro G. Salerno, Shathiyah Kulandavelu, Wayne Balkan, Marilia S.S.R. Zuttion, Renzhi Cai, Andrew V. Schally, Joshua M. Hare

## Abstract

**Objective:** To test the hypothesis that the activation of the growth hormone-releasing hormone (GHRH) receptor signaling pathway within the myocardium both prevents and reverses heart failure with preserved ejection fraction (HFpEF).

**Background:** HFpEF is characterized by impaired myocardial relaxation, fibrosis and ventricular stiffness. Despite the rapidly increasing prevalence of HFpEF, no effective therapies have emerged. Synthetic agonists of the GHRH receptors reduce myocardial fibrosis, hypertrophy and improve performance, independently of the growth-hormone axis.

**Methods:** We generated a HFpEF-like phenotype with continuous infusion of angiotensin-II (Ang-II) in CD1 mice. Mice were injected with either vehicle or a potent synthetic agonist of the growth hormone-releasing hormone, MR-356.

**Results:** Ang-II treated animals had diastolic dysfunction, ventricular hypertrophy, and normal ejection fraction and isolated cardiomyocytes (ex vivo) exhibited incomplete relaxation, depressed contractile responses and altered myofibrillar protein phosphorylation. Calcium handling mechanisms were disturbed in cardiomyocytes from mice with HFpEF. MR-356 both prevented and reversed the development of the pathological phenotype in vivo and ex vivo.

**Conclusion:** These findings indicate that the GHRH receptor signaling pathway represents a new molecular target to counteract HFpEF-associated cardiomyocyte dysfunction by targeting myofilament phosphorylation. Accordingly, activation of the GHRH receptor with potent synthetic GHRH agonists may provide a novel therapeutic approach to management of the HFpEF syndrome.

**Condensed abstract:** Heart failure with preserved ejection fraction (HFpEF) is characterized by a remodeled myocardium conferring ventricular stiffness and diastolic dysfunction. There are no effective therapies. Agonists of growth hormone-releasing hormone (GHRH) receptors have beneficial effects on the heart. We hypothesize that activation of GHRH receptors suppresses this HFpEF phenotype. Treatment with a synthetic agonist of GHRH, prevented the development of the pathological phenotype in a murine model of HFpEF-induced by chronic angiotensin-II infusion. These findings indicate that activation of GHRH receptors represents a novel molecular strategy to counteract HFpEF-associated cardiomyocyte dysfunction and provide a potential approach to management of HFpEF syndrome.

**Highlights:**

- A synthetic growth hormone-releasing hormone agonist (GHRH-A) prevents and reverses the pathological remodeling in a mouse model of HFpEF induced by infusion of low dose Ang II.
- GHRH-A improves intracellular calcium handling by reducing the sarcoplasmic reticulum calcium leakage and enhancing phospholamban phosphorylation.
- GHRH-A treatment prevents and reverses diastolic dysfunction by enhancing the rate of sarcomere re-lengthening.
- Activation of the GHRH receptor with the GHRH-A, MR-356, leads to targeting myofibrillar proteins and desensitizing myofilaments in response to calcium.

## Introduction

Heart failure (HF) remains a leading cause of death worldwide and is associated with two general clinical variants – heart failure with reduced ejection fraction (HFrEF), characterized by systolic dysfunction, and heart failure with preserved ejection fraction (HFpEF), predominantly due to diastolic dysfunction(1–3). HFpEF accounts for ~50% of HF cases, a proportion that is predicted to increase yearly relative to HFrEF(4). Numerous drugs treat HFrEF; no approved therapies target HFpEF, and clinical trials with several classes of drugs, including renin-angiotensin-aldosterone inhibitors, produced no clinical benefits(5–8). The HFpEF syndrome is characterized by endothelial dysfunction, hypertension, impaired relaxation and contractile reserve, ventricular stiffening and fibrosis(2,9). The underlying molecular mechanisms are poorly understood due to diverse etiology, generally involving multi-organ pathological conditions(10–12). Importantly, current therapeutic strategies fail to abrogate the progression of HFpEF(5,7,13,14), thereby creating a healthcare crisis(15,16).

The development of new potent synthetic growth hormone-releasing hormone agonists (GHRH-As) and their application in different models of myocardial injury (ischemic(17–21) or non-ischemic(22)) demonstrates that activation of myocardial GHRH receptors (GHRHRs) effectively reduces fibrosis or hypertrophy, respectively, and improves cardiac performance(18–22). GHRHR signaling extends beyond the hypothalamic-pituitary circuit(23). GHRHRs are present on cardiomyocytes(18–20), and GHRH-As signal in the myocardium acts independently of the growth hormone (GH)/insulin-like growth factor 1(IGF1) axis(19).

Few animal models reproduce features of HFpEF(11,24–26). We used chronic infusion of low dose angiotensin II (Ang-II) in mice, which mimics the HFpEF cardiac phenotype(27), to study the effects of a synthetic GHRH-A, MR-356, on cardiomyocyte performance *in vivo* and *ex vivo*. MR-356 is ~150x more potent(17) than human GHRH(1–29)NH_2_ and binds the GHRHR with 7.5-fold greater affinity. Therefore, we hypothesized that GHRH-As improve cardiomyocyte function in HFpEF and that activation of this GHRHR signaling both prevents and reverses features of myocardial HFpEF by targeting both impaired cardiomyocyte relaxation and myocardial fibrosis.

## Methods

### Experimental design and animal model

Male CD1 mice (Envigo) were implanted with a mini-osmotic pump (Alzet) to deliver Ang-II (0.2 mg/kg/day; Sigma-Aldrich Co., Saint Louis, MO) for 4 weeks. One set of mice received daily injections of GHRH-Agonist (GHRH-A [MR-356]: 200 μg/kg) or vehicle (DMSO+propylene-glycol) for 4 weeks. during the same period. The second group of mice received only the mini-pump, which was replaced for an additional 4-week Ang-II infusion, at which point mice were randomized to receive 4-weeks of daily injection of MR-356 or vehicle. Unmanipulated age-matched CD1 mice acted as normal controls. All protocols and experimental procedures were approved by the University of Miami Animal Care and Use Committee and followed the Guide for the Care and Use of Laboratory Animals (NIH Publication No.85-234, revised 2011).

### Materials

MR-356 ([N-Me-Tyr^1^, Gln^8^, Orn^12^, Abu^15^, Orn^21^, Nle^27^, Asp^28^, Agm^29^] hGH-RH(1–29)), was synthetized by solid phase methods and purified by HPLC in our laboratories(17).

### Echocardiography

Transthoracic echocardiography was performed as previously described with minor modifications(28). All echocardiographic measurements were performed offline over 3 to 5 heart beats.

### Speckle Tracking Echocardiography (STE)

Strain analysis was conducted using speckle-tracking software, Vevostrain TM analysis (Visual Sonics). Global longitudinal strain was calculated using B-mode cine images in the LV parasternal long-axis view acquired at a high frame rate(29).

### Hemodynamic Measurements

Hemodynamic studies were performed using a micro-tipped pressure-volume catheter (SPR-839; ADInstruments) as described(30) with minor modifications. All analyses were performed using LabChart Pro version 8.1.5 software (ADInstruments)

### Cardiomyocyte Isolation

Cardiomyocytes were isolated and prepared from hearts of mice by retrograde perfusion through the aorta in a modified Langerdorf system with collagenase type 2 and protease type XIV as described previously(31).

### Intracellular Calcium and Sarcomere Length Measurement

Intracellular Ca^2+^ was measured using Fura-2 and a dual-excitation (340/380 nm) spectrofluorometer (IonOptix LLC) in cardiomyocytes electric-field stimulated at 1, 2 and 4 Hz. Calcium concentration ([Ca^2+^]_i_) was calculated as described(31). Simultaneously, sarcomere length (SL) was recorded with an IonOptix iCCD camera.

Hysteresis loops were obtained by plotting sarcomere length versus [Ca^2+^]_i_ of cardiomyocytes paced at 4 Hz.

SRCa^2+^ leakage was assessed with 1 mmol/L tetracaine (Sigma-Aldrich) as described by Shannon *et al*.(32) with some modifications. SR Ca^2+^ content was assessed on the same cell by 20 mmol/L caffeine pulse as described previously(31). Measurements were carried out at 37°C.

### Western Blot Immunoanalysis

Protein expression of SERCA2a, PLB, L-TCC, NCX1, cTnI, cMyBPC, collagen I and III, TGF-β, P4HA1 and protein phosphorylation analyses were assessed by Western blot. Briefly, supernatants from isolated cardiomyocyte or heart tissue homogenates were collected and protein concentration determined. Samples were electrophoresed, transferred to PVDF or nitrocellulose membranes (Bio-Rad Laboratories) and detected by immunoblotting as indicated.

### Interstitial Fibrosis Quantification

Paraffin-fixed heart sections were subjected to Masson’s trichrome staining for assessment of fibrillary collagen deposition. Images were acquired by an Olympus VS120 scanner and analyzed with ImageJ.

### Quantitative RT-PCR

Total RNA was isolated from cardiomyocytes using RNAeasy mini kit (Qiagen, Hilden, Germany). RNA (1 μg) was reverse-transcribed (Applied Biosystems, Foster City, CA) using Taqman^®^ Universal Master Mix (Applied Biosystems, Foster City, CA) in a Bio-Rad CFX96™ Real-Time PCR system (Bio-Rad Laboratories, Hercules, CA, USA). Change in mRNA expression was normalized to the change in 18S, and values were expressed as 2^ΔCt^.

Detailed methodology is provided in the Supplemental Material.

### Statistical Analysis

Data are reported as mean ± standard error of the mean (SEM). Statistical significance between three groups was determined by one-way or two-way ANOVA followed by Tukey’s or Bonferonni’s post hoc tests, as appropriate. For comparisons of two groups, Student’s two-tailed t-test was used. Analyses were performed using GraphPad Prism version 8.4.3. The null hypothesis was rejected at *p*<0.05.

## Results

### GHRH-A prevents the appearance of the HFpEF syndrome induced by in vivo Ang-II infusion

Four-weeks of continuous infusion of Ang-II induced a cardiovascular phenotype that mimicked HFpEF phenotypically(27), specifically, myocardial hypertrophy with increased heart weight normalized by body weight or tibia length (Figure 1A and Supplemental Table 1), increased relative wall thickness in diastole (RWTd) and increased cardiomyocyte width (Figure 1B and Table 1, respectively) without changes in length (Supplemental Figure 1). Myocardial fibrosis increased in the HFpEF hearts as indicated by collagen III content and the transforming growth factor-β (TGF-β), a fibrosis mediator (Figure 1C-D). Prolyl-4-hydroxylase-α-1 (P4HA1), and the hydroxyproline residues were slightly increased while collagen I did not change (Supplemental Figure 2A-C). Co-administration of the GHRH-A, MR-356, prevented the increases in cardiomyocyte width and RWTd (Figure 1B and Table 1).Moreover, fibrosis was also prevented by MR-356 since collagen III was reduced and TGF-β levels controlled (p=0.234 versus control)(Figure 1C-D).

**Figure 1:**
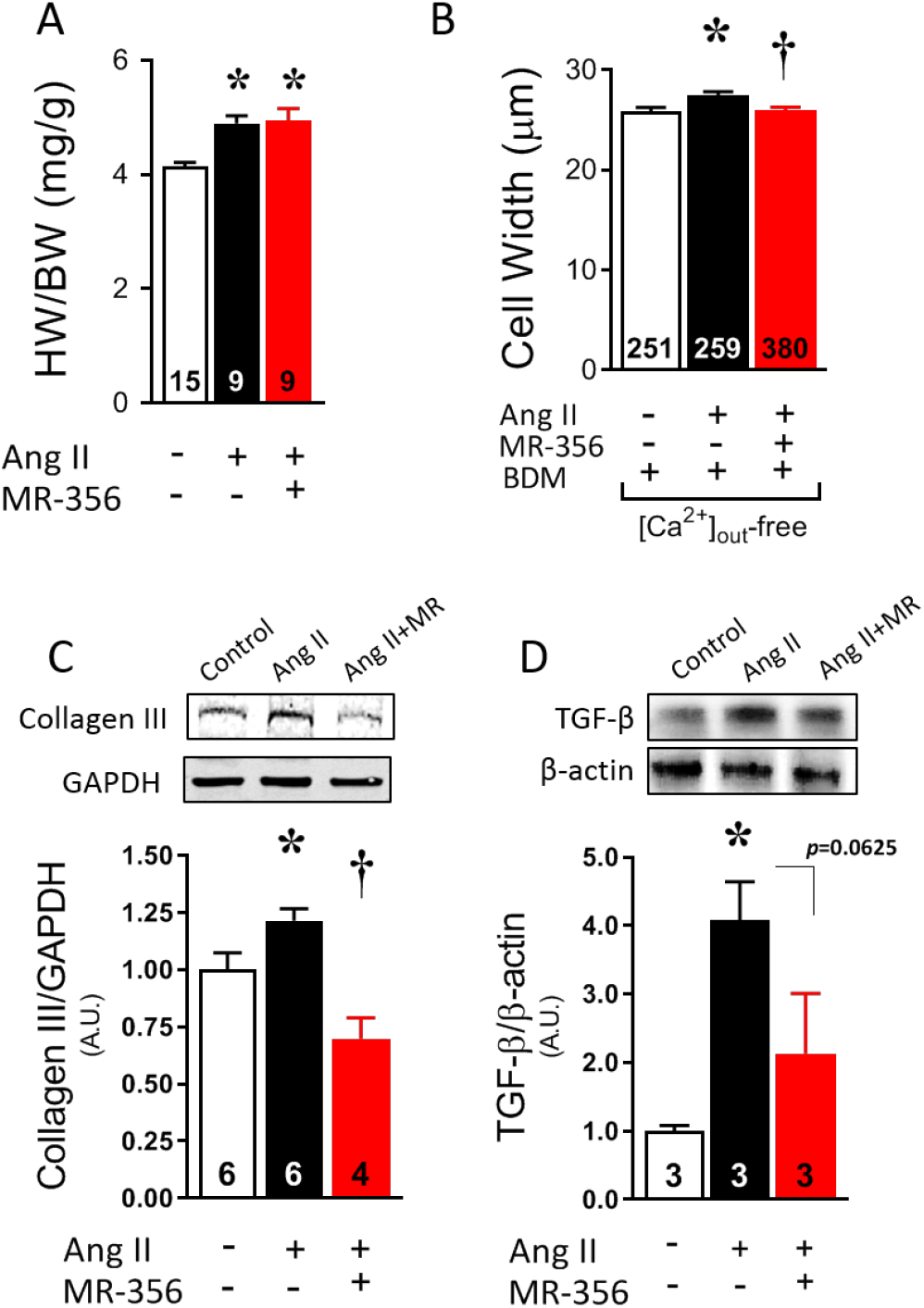
GHRH-A MR-356 prevents cardiomyocyte growth and fibrosis. (A) Heart weight normalized to body weight. (B) Width of isolated cardiomyocytes from control (N=4), Ang-II-treated (N=5) and Ang-II+MR-356-treated (N=5) mice. (C) Collagen III protein expression in heart homogenates. Top, representative images of Western blots. Bottom, average results. (D) TGF-β protein expression in cardiomyocytes. \**p*<0.05 in Ang II vs. control and † *p*<0.05 in Ang II+MR-356-treated vs. Ang II; One-way ANOVA. Numbers at bottom of bars correspond to studied animals in panel A, number of cells in panel B, and number of samples in panels C-D.

**Table 1:**
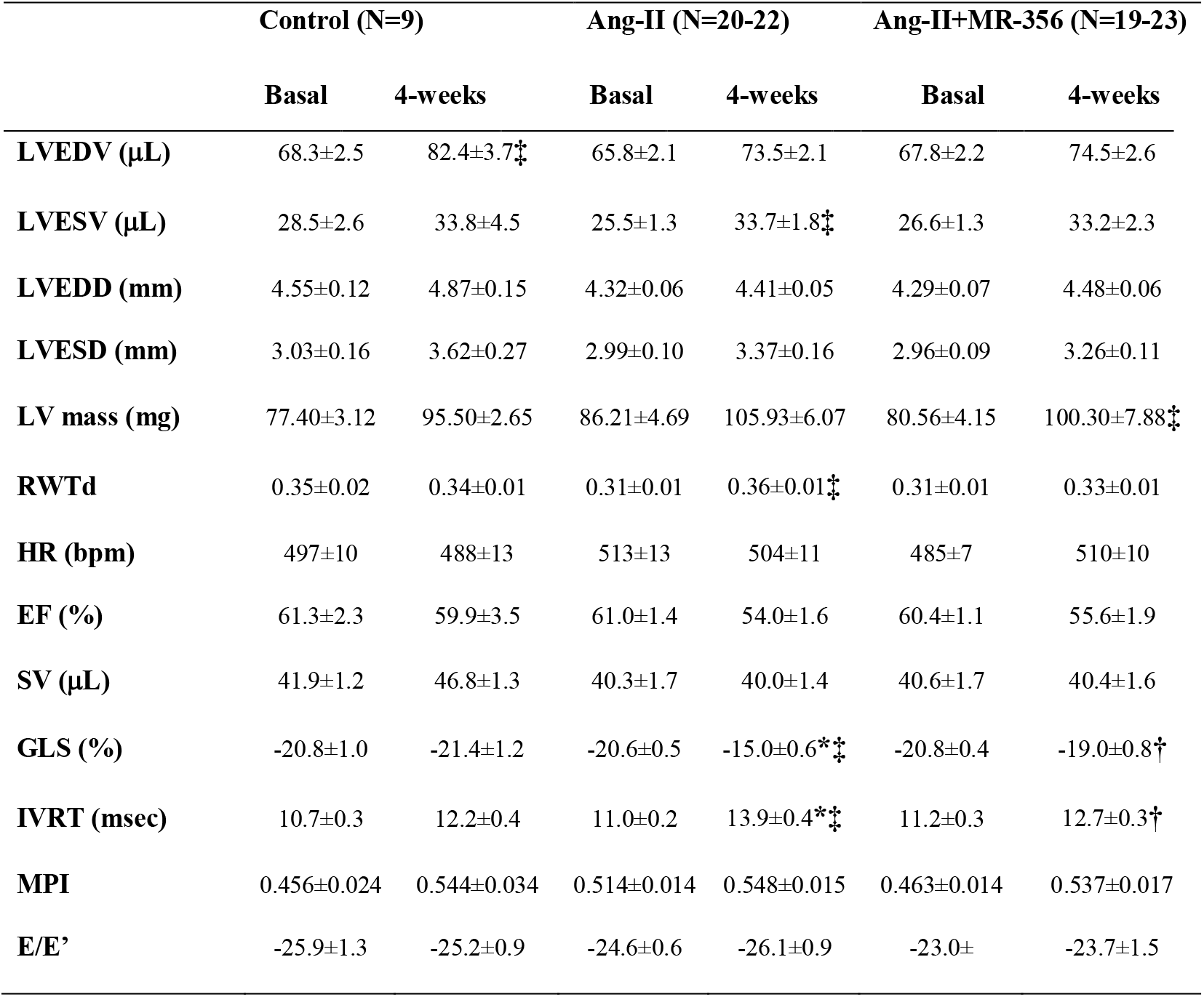
Echocardiographic parameters of systolic and diastolic function in 4-week Ang-II infusion model. All values represent mean±SEM. ‡*p*<0.05 at 4-week vs. baseline; \**p*<0.05 vs. control; †*p*<0.05 Ang-II+MR-356 vs. Ang-II, two-way ANOVA. N represents number of studied animals.

|  | Control (N=9) |  | Ang-II (N=20-22) |  | Ang-II+MR-356 (N=19-23) |  |
| --- | --- | --- | --- | --- | --- | --- |
|  | Basal | 4-weeks | Basal | 4-weeks | Basal | 4-weeks |
| **LVEDV (μL)** | 68.3±2.5 | 82.4±3.7‡ | 65.8±2.1 | 73.5±2.1 | 67.8±2.2 | 74.5±2.6 |
| **LVESV (μL)** | 28.5±2.6 | 33.8±4.5 | 25.5±1.3 | 33.7±1.8‡ | 26.6±1.3 | 33.2±2.3 |
| **LVEDD (mm)** | 4.55±0.12 | 4.87±0.15 | 4.32±0.06 | 4.41±0.05 | 4.29±0.07 | 4.48±0.06 |
| **LVESD (mm)** | 3.03±0.16 | 3.62±0.27 | 2.99±0.10 | 3.37±0.16 | 2.96±0.09 | 3.26±0.11 |
| **LV mass (mg)** | 77.40±3.12 | 95.50±2.65 | 86.21±4.69 | 105.93±6.07 | 80.56±4.15 | 100.30±7.88‡ |
| **RWTd** | 0.35±0.02 | 0.34±0.01 | 0.31±0.01 | 0.36±0.01‡ | 0.31±0.01 | 0.33±0.01 |
| **HR (bpm)** | 497±10 | 488±13 | 513±13 | 504±11 | 485±7 | 510±10 |
| **EF (%)** | 61.3±2.3 | 59.9±3.5 | 61.0±1.4 | 54.0±1.6 | 60.4±1.1 | 55.6±1.9 |
| **SV (μL)** | 41.9±1.2 | 46.8±1.3 | 40.3±1.7 | 40.0±1.4 | 40.6±1.7 | 40.4±1.6 |
| **GLS (%)** | -20.8±1.0 | -21.4±1.2 | -20.6±0.5 | -15.0±0.6*‡ | -20.8±0.4 | -19.0±0.8† |
| **IVRT (msec)** | 10.7±0.3 | 12.2±0.4 | 11.0±0.2 | 13.9±0.4*‡ | 11.2±0.3 | 12.7±0.3† |
| **MPI** | 0.456±0.024 | 0.544±0.034 | 0.514±0.014 | 0.548±0.015 | 0.463±0.014 | 0.537±0.017 |
| **E/E'** | -25.9±1.3 | -25.2±0.9 | -24.6±0.6 | -26.1±0.9 | -23.0± | -23.7±1.5 |

Echocardiographic evaluation demonstrated that Ang-II infusion did not reduce ejection fraction (EF) (Table 1), either alone or under co-treatment with MR-356. Likewise, chamber dimensions were unaffected in systole between groups (Table 1). Invasive hemodynamic assessment (Table 2) demonstrated increased arterial blood pressure and clear evidence of diastolic dysfunction with increased end-diastolic pressure-volume relationship (EDPVR) (Figure 2A-D), relaxation time constant (tau) and left ventricular end-diastolic pressure (LVEDP) in Ang-II-treated mice compared to control (Figure 2E-F). Isovolumic relaxation time (IVRT) increased along the time (Figure 2H) (within group comparison - 4-week versus baseline), and this increase was greater in the Ang-II-treated. Global longitudinal strain (GLS) increased (reduced absolute values) with Ang-II treatment, in an extent according to that observed in diastolic dysfunction(33) (Figure 2I and Table 1). Co-infusion of MR-356 completely prevented the development of diastolic dysfunction (Figure 2C-I) and promoted a robust acceleration in dP/dt_*min*_ (Figure 2G). Together, these results demonstrate that stimulation of GHRH pathway with MR-356 effectively counteracted the development of Ang-II-induced HFpEF.

**Figure 2:**
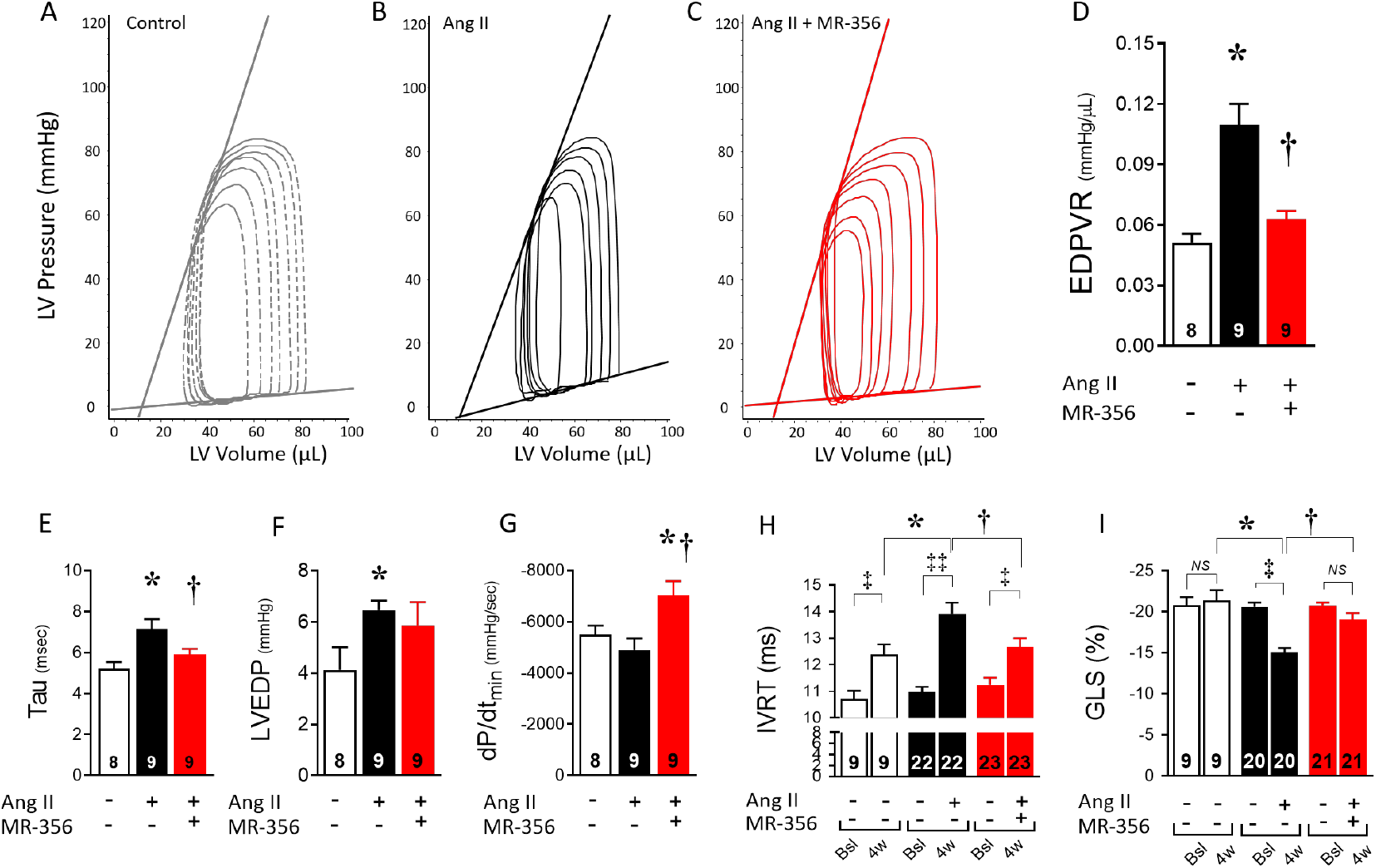
Effects of the GHRH-A, MR-356, on diastolic dysfunction in Ang-II-induced HFpEF mice. (A) Representative pressure-volume (P-V) loops showing the linear regression for ESPVR and EDPVR in a control, (B) Ang-II-treated, or (C) Ang-II+MR-356 mouse. (D) Averaged slopes from the EDPVR linear fitting. (E) Hemodynamic assessment of the relaxation time constant, tau, at 4-week time point. (F) LVEDP. (G) Maximal rate of pressure decline (*d*P/*dt*_min_). (H) IVRT as measured by transthoracic echocardiography at baseline or after 4-week treatment. (I) GLS analysis of the left ventricle. GLS represents the function of sub-endocardial longitudinal myocardial fibers that are more sensitive to reduced coronary perfusion and increased wall stress. Moreover, GLS reflects changes in the myocardial interstitium including the extent of myocardial fibrosis. ^‡^*p*<0.05 at 4-week vs. baseline; ^‡‡^*p*<0.001 at 4-week vs. baseline, two-way ANOVA. \**p*<0.05 vs. control; ^†^*p*<0.05 in Ang-II+MR-356-treated vs. Ang-II; one- or two-way ANOVA. Numbers on bottom of bars indicate number of analyzed animals.

**Table 2:** Hemodynamic assessments on 4-week Ang-II infusion model. All values represent mean±SEM. \**p*<0.05 vs. control; †*p*<0.05 Ang-II+MR-356 vs. Ang-II, one-way ANOVA. N represents number of studied animals.

|  | Control (N=8) | Ang-II (N=9) | Ang-II+MR-356 (N=9) |
| --- | --- | --- | --- |
| HR (1/min) | 535.9±11.7 | 543.9±7.2 | 523.5±14.2 |
| EF (%) | 51.0±4.0 | 55.7±5.8 | 59.6±2.9 |
| Tau (1/sec) | 5.21±0.33 | 7.12±0.57* | 5.91±0.27† |
| dP/dt<sub>max</sub> (mmHg/sec) | 7328±346 | 6866±519 | 7433±476 |
| dP/dt<sub>min</sub> (mmHg/sec) | -5494±353 | -4897±451 | -7013±580*† |
| Ea (mmHg/μL) | 1.88±0.25 | 1.85±0.29 | 1.94±0.13 |
| LVEDP (mmHg) | 4.13±0.89 | 6.46±0.38* | 5.87±0.91 |
| ESPVR (mmHg/μL) | 2.71±0.28 | 1.85±0.32* | 2.03±0.21* |
| EDPVR (mmHg/μL) [linear] | 0.051±0.004 | 0.110±0.010* | 0.063±0.004† |
| PRSW (mmHg/μL) | 74.5±3.1 | 63.2±3.9 | 62.4±5.4 |
| dP/dt<sub>max</sub>-EDV (mmHg/μL.sec) | 110.7±13.9 | 99.0±6.3 | 91.1±9.6 |
| Ea/Ees | 0.72±0.05 | 1.15±0.15* | 1.01±0.11 |
| Systolic ABP (mmHg) | 79.7±2.2 | 103.0±4.7* | 88.5±4.6† |
| Diastolic ABP (mmHg) | 54.7±2.7 | 75.5±3.7* | 64.4±4.2 |

### GHRH-A prevents sarcomere and contractile dysfunction in HFpEF cardiomyocytes

To test whether GHRH agonists directly affected cardiomyocyte function, isolated cardiomyocytes were electric field-stimulated at 1, 2 and 4 Hz. Cardiomyocytes from Ang-II-infused mice exhibited shorter resting (diastolic) sarcomere length compared to those from control animals (Figure 3A, B), indicating an inability to relax. In parallel, quiescent HFpEF cardiomyocytes exposed to 1.8 mmol/L extracellular CaCl_2_ were consistently shorter than control cardiomyocytes, while there was no difference between groups in Ca^2+^-free buffer in the presence of 2,3-butanedione monoxime (BDM) (Supplemental Figure 3A). HFpEF cardiomyocytes also exhibited depressed contractile response to pacing (Figure 3A, C). However, MR-356 administration during Ang-II infusion, both prevented impairment of resting sarcomere tension (Figure 3A, B), and preserved quiescent cardiomyocyte length in the presence of extracellular Ca^2+^ (Supplemental Figure 3A, right panel). Treatment with MR-356 also preserved sarcomere shortening and contractile response to pacing as in control cardiomyocytes (Figure 3A, C). There were no significant differences in Ca^2+^ transient amplitude (Δ[Ca^2+^]) between groups (Figure 3D, E), which suggests an altered responsiveness to Ca^2+^. However, when Δ[Ca^2+^] was normalized, an impaired response to pacing was observed in HFpEF cardiomyocytes, while the MR-356-treated ones conserved the positive response to pacing as controls (Figure 3F). This finding correlates with their contractile response (Figure 3C).

**Figure 3:**
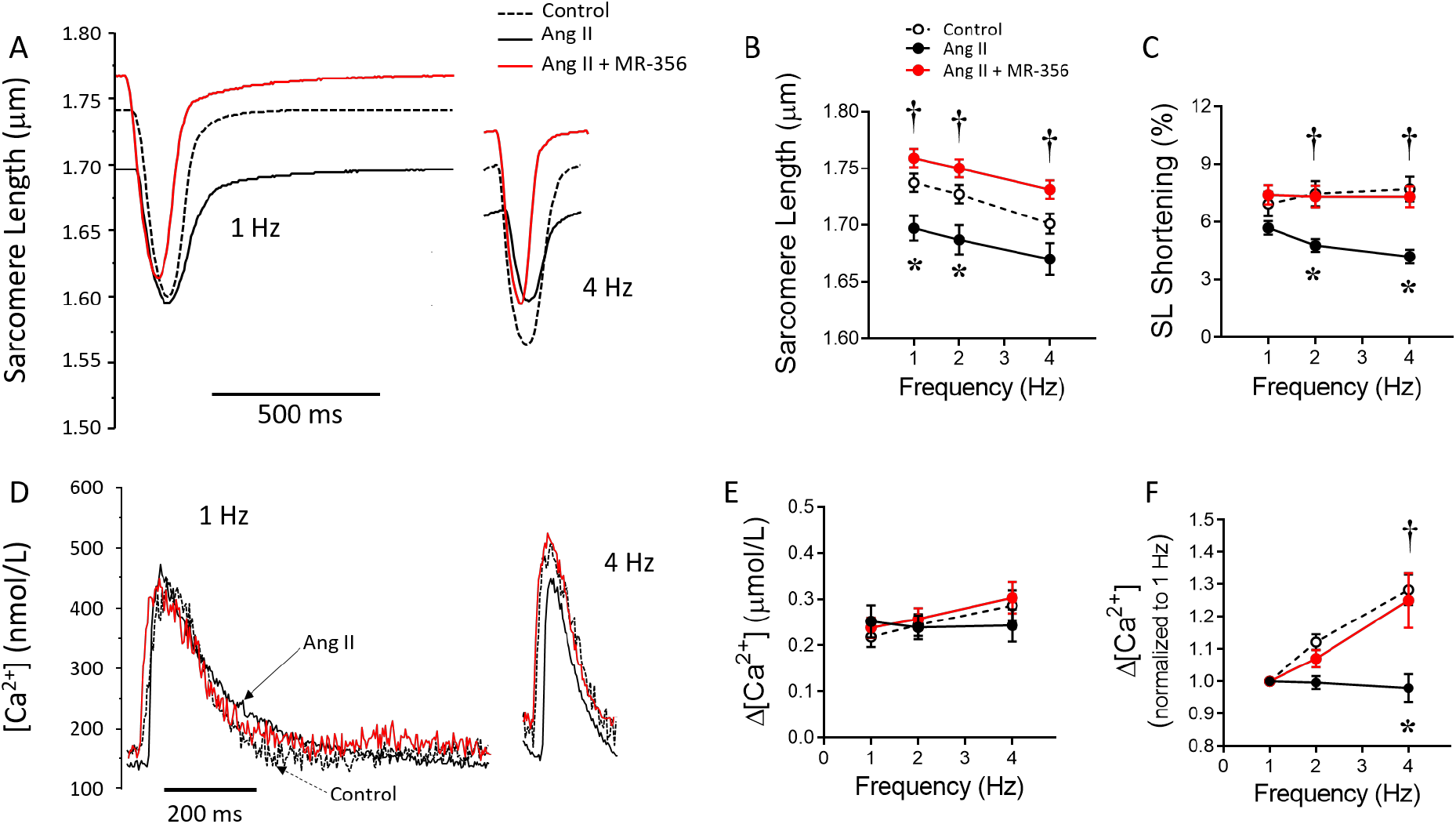
Sarcomere and contractile dysfunction in the HFpEF model is prevented by co-treatment with GHRH-A MR-356. (A) Representative traces of sarcomere twitches of cardiomyocytes, electric field-stimulated at 1- and 4-Hz, from male CD1 control mice (N=4, 30 cells) or mice that underwent either Ang-II (N=5, 41 cells) or Ang-II+MR-356 treatment (N=5, 38 cells). (B) Resting sarcomere length in cardiomyocytes from each group at different pacing rates. (C) Sarcomere shortening in response to pacing. (D) Representative traces of intracellular [Ca^2+^] in electric field-stimulated cardiomyocytes. (E) Δ[Ca^2+^] transient amplitude in response to pacing. (F) Δ[Ca^2+^] normalized (respect to 1 Hz) in response to pacing. From each mouse, a total of 5-9 cells were studied. \**p*<0.05 in Ang-II vs. control and ^†^*p*<0.05 in Ang-II+MR-356-treated vs. Ang-II; two-way ANOVA.

Increased SR Ca^2+^ leak is a hallmark of heart failure and contribute to depressed contractile reserve. We assessed sarcoplasmic reticulum (SR) Ca^2+^ leak with tetracaine (see Supplemental Material and Figure 4A). We found that Ang-II-treated myocytes exhibited a steeper rise in SRCa^2+^leak with increasing SRCa^2+^ loads compared with controls (*p*<0.05) and this load-leak curve was flattened by treatment with MR-356 (*p*<0.0001; Figure 4B). Thus, at matched SRCa^2+^ load=99.1±11.5 μmol/L, the Ca^2+^ leak was elevated in Ang-II-induced HFpEF cardiomyocytes and partially prevented by treatment with MR-356 (Figure 4C).

**Figure 4:**
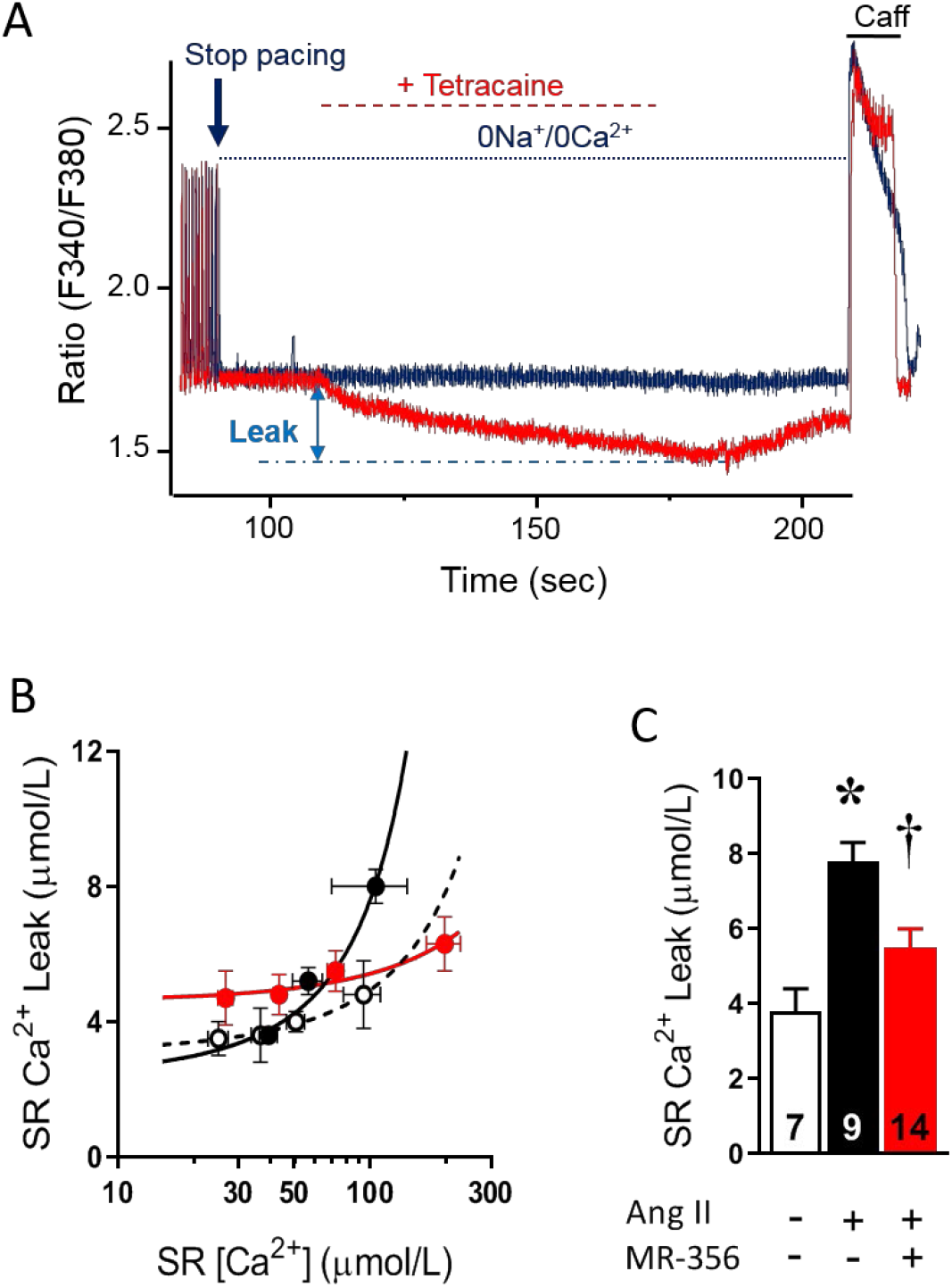
Increased SRCa^2+^ leak in HFpEF cardiomyocytes is prevented by treatment with the GHRH-A MR-356. (A) Representative traces of intracellular [Ca^2+^] in cardiomyocytes illustrating the method for measuring SRCa^2+^ leak using tetracaine. Blue trace corresponds to Ca^2+^ from a cardiomyocyte not exposed to tetracaine; red trace is the Ca^2+^ from the same cardiomyocyte treated with tetracaine for 70 seconds after pacing was stopped. (B) SRCa^2+^ leak-load relationship in cardiomyocytes from control (n=31 cells), Ang-II-treated (n=38 cells) or Ang-II+MR-356-treated (n=35 cells) CD1 mice. (C) Average SRCa^2+^ leak in cardiomyocytes from all groups at matched SRCa^2+^ load=99.1 μmol/L. \**p*<0.05 in Ang-II vs. control and ^†^*p*<0.05 in Ang-II-treated vs. Ang-II+MR-356-treated; one- or two-way ANOVA as appropriate. Numbers on bottom of bars indicate number of analyzed cells.

### Effect of GHRH-A on the impaired Ca^2+^ decline in HFpEF cardiomyocytes

We found slower [Ca^2+^] decay in HFpEF cardiomyocytes compared with controls and treatment with MR-356 blocked the delayed Ca^2+^ decline (Figure 5A, B). The Ca^2+^ content in the SR, estimated by a rapid caffeine pulse, was not different from control. Ang-II+MR-356 co-treatment did increase the SR [Ca^2+^] in comparison to control cardiomyocytes (Figure 5C). We investigated the proteins participating in Ca^2+^ handling. The SR Ca^2+^ ATPase (SERCA2) protein expression was reduced in cardiomyocytes from Ang-II-treated mice (Figure 5D). Phospholamban (PLB) phosphorylation at serine 16 (p-Ser16) or threonine 17 (p-Thr17) were increased by Ang-II treatment, an effect further enhanced by co-treatment with MR-356 (Figure 5E, F).

**Figure 5:**
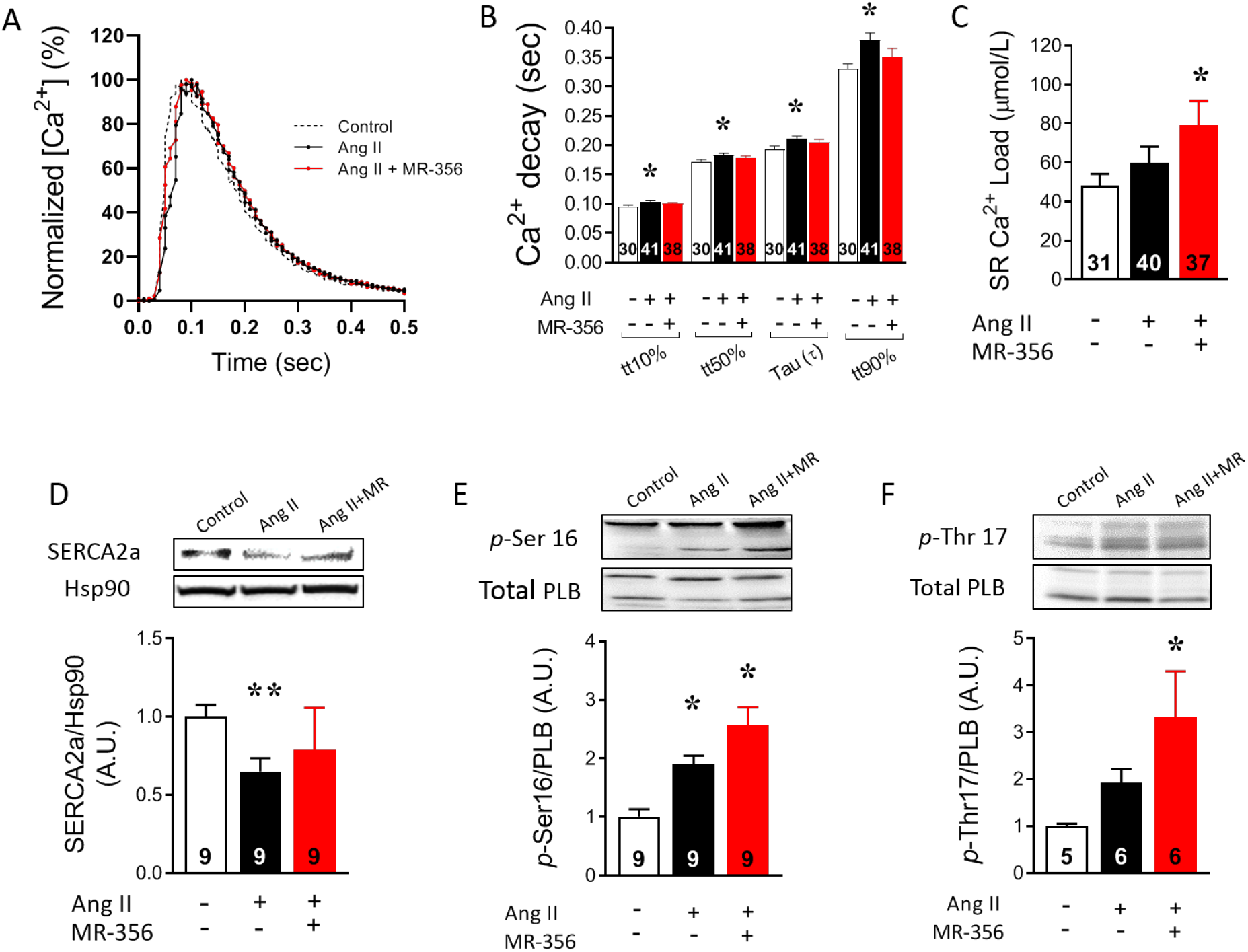
Ca^2+^ decay is delayed in cardiomyocytes from the Ang-II-induced HFpEF model but preserved by treatment with MR-356. (A) Normalized Ca^2+^ transient re-constructed by using average dynamic parameters from control untreated, Ang-II-treated or Ang-II+MR-356-treated cardiomyocytes paced at 1 Hz. (B) Average times to different degrees of the Ca^2+^ peak decline: time to 10% (tt10%), time to 50% (tt50%), decay time constant (τ) and time to 90% (tt90%). (C) SR Ca^2+^ content in cardiomyocytes paced at 1 Hz. Numbers on bottom of bars on panels B and C indicate number of analyzed cells. (D) SERCA2a protein expression; top, representative immunoblotting image with Hsp90 as loading control and bottom, average normalized values. (E) PLB phosphorylation at serine 16. (F) Phosphorylation of PLB at threonine 17. Top, representative images and bottom, average values normalized by total PLB expression. Numbers on bottom of bars on panels D, E and F indicate number of analyzed samples. \**p*<0.05 or \*\**p*<0.01 vs. control, ^†^*p*<0.05 vs. Ang-II-treated; one-way ANOVA.

### Impaired cardiomyocyte relaxation in HFpEF is improved by GHRH-A

Myocardial stiffness and deficient relaxation are hallmarks of the HFpEF syndrome, and may be due to factors external to the cardiomyocyte or intrinsically associated with sarcomeric proteins. At the cardiomyocyte level, this mouse model exhibited reduced ability to relax, mainly during the late part of the relaxation phase (tau and tt90%, Fig 6A, B). Treatment with MR-356 dramatically improved all phases of sarcomere relaxation (Figure 6A, B). Comparing this effect of MR-356 with the modest effect on Ca^2+^ decline rate, suggests a direct targeting on myofilaments. Accordingly, we analyzed SL-[Ca^2+^] hysteresis loops of cardiomyocyte contractions induced by electric-field stimulation, in order to detect differences in responsiveness to Ca^2+^. Overall, loop morphology was greatly affected in the HFpEF model (Figure 6C). In the relaxation phases (Figure 6C, D left), there was Ang-II-induced sensitization of myofilaments (*p*<0.0001, Figure 6D right) consistent with the delayed Ca^2+^ decline in this model. Co-treatment with MR-356 induced a robust desensitizing effect that correlated with the exacerbated lusitropic effect of MR-356 on isolated cardiomyocytes (Figure 6D). We explored phosphorylation on sites of cardiac troponin I (cTnI) with regulatory activity, serine 23/24. TnI phosphorylation was slightly impaired in Ang-II-treated cardiomyocytes (p=0.071) while co-treatment with MR-356 promoted increased phosphorylation (Figure 6E). Similarly, HFpEF cardiomyocytes exhibited reduced phosphorylation at serine 282 of the myosin binding protein C (MyBPC), which was prevented in Ang-II+MR-356 cardiomyocytes (Figure 6F). These findings consistently correlate with the data of responsiveness to Ca^2+^ (Figure 6D).

**Figure 6:**
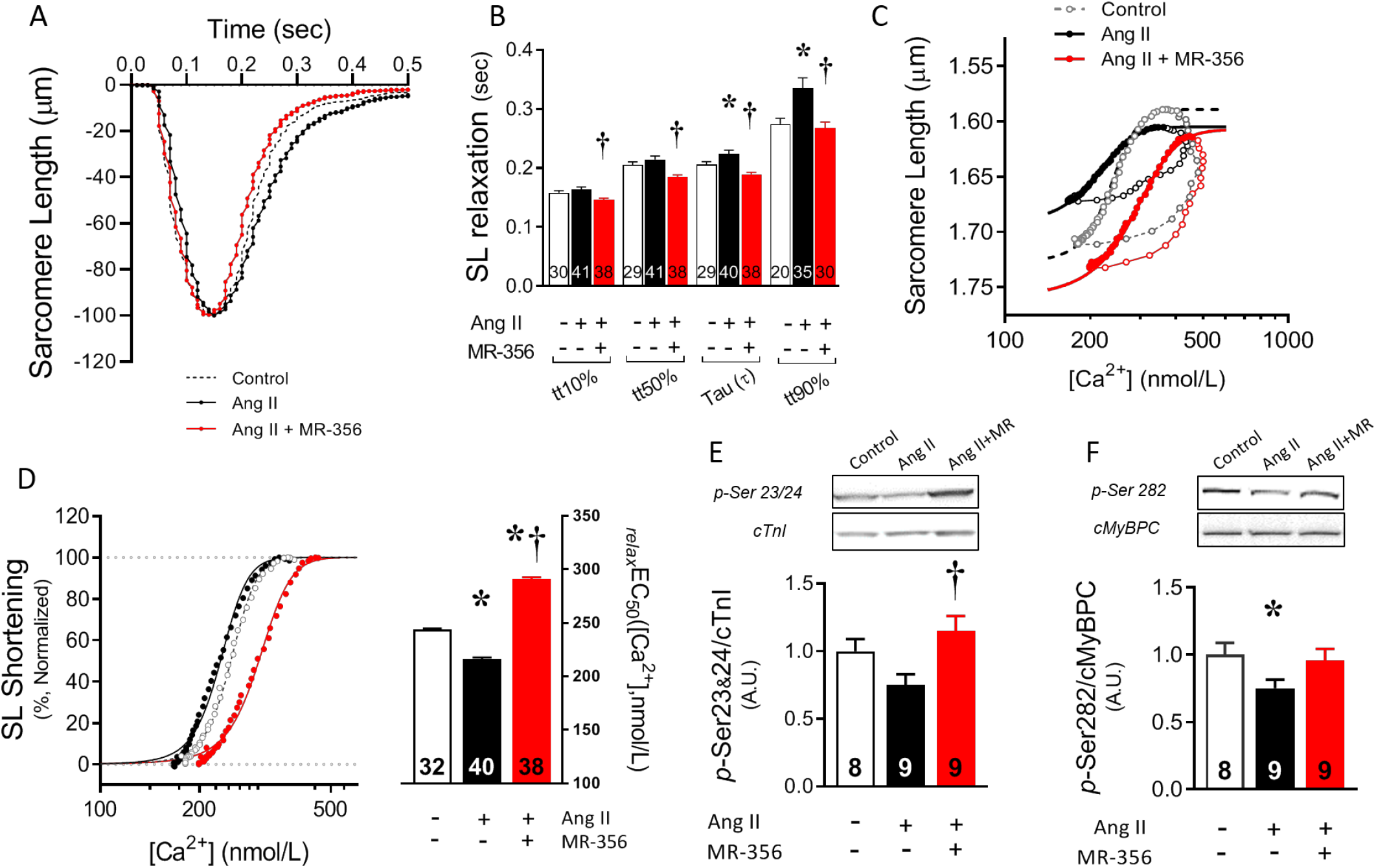
Impaired sarcomere relaxation due to myofilament sensitization in HFpEF cardiomyocytes is prevented by treatment with MR-356. (A) Normalized sarcomere contraction re-constructed by using average SL dynamic parameters at 1Hz from control untreated, Ang-II-treated or Ang-II+MR-356-treated cardiomyocytes. (B) Average times to different degrees of the relaxation: tt10%, tt50%, tau (τ) and tt90%. (C) SL-[Ca^2+^] hysteresis loops in control, Ang-II-or Ang-II+MR-356-treated cardiomyocytes with overlapped sigmoidal fitting of the relaxation phases. (D) Normalized relaxation phase of SL-[Ca^2+^] loops (left) and averaged effective [Ca^2+^] of the relaxation phase (_*relax*_EC_50_s, right). Numbers on bottom of bars in panel D indicate number of analyzed cells from 4-5 hearts. (E) Phosphorylation of cTnI at serine 23/24 and total cTnI protein expression by western blot. (F) Phosphorylation of cMyBPC at serine 282 and total cMyBPC protein. Top, representative images; bottom, average normalized values. Numbers on bottom of bars on panels E and F indicate number of analyzed samples. \**p*<0.05 vs. control, ^†^*p*<0.05 vs. Ang-II + MR-356-treated; one-way ANOVA.

### GHRH-A improves relaxation and reverts the HFpEF phenotype induced by eight-week Ang-II infusion in CD1 mice

Given the ability of GHRH-As to prevent the onset of HFpEF, we next performed experiments to test whether GHRH-As reverse established HFpEF. We infused mice with a low dose of Ang-II for 8 weeks and initiated daily treatment with either MR-356 or vehicle beginning at week 4, once the HFpEF-like phenotype was already apparent. As with the 4-week protocol, myocardiac hypertrophy (increased LV mass) induced by Ang-II, was not significantly reduced by MR-356 treatment (Figure 7A). However, the molecular remodeling (as indicated by the altered ratio between the adult (α) to fetal (β) isoform of the myosin heavy chain gene expression, MHY6/MHY7) was normalized by MR-356 treatment (Figure 7B). The increased interstitial fibrosis in HFpEF hearts was consistently reverted by MR-356 (Figure 7C-F).

**Figure 7:**
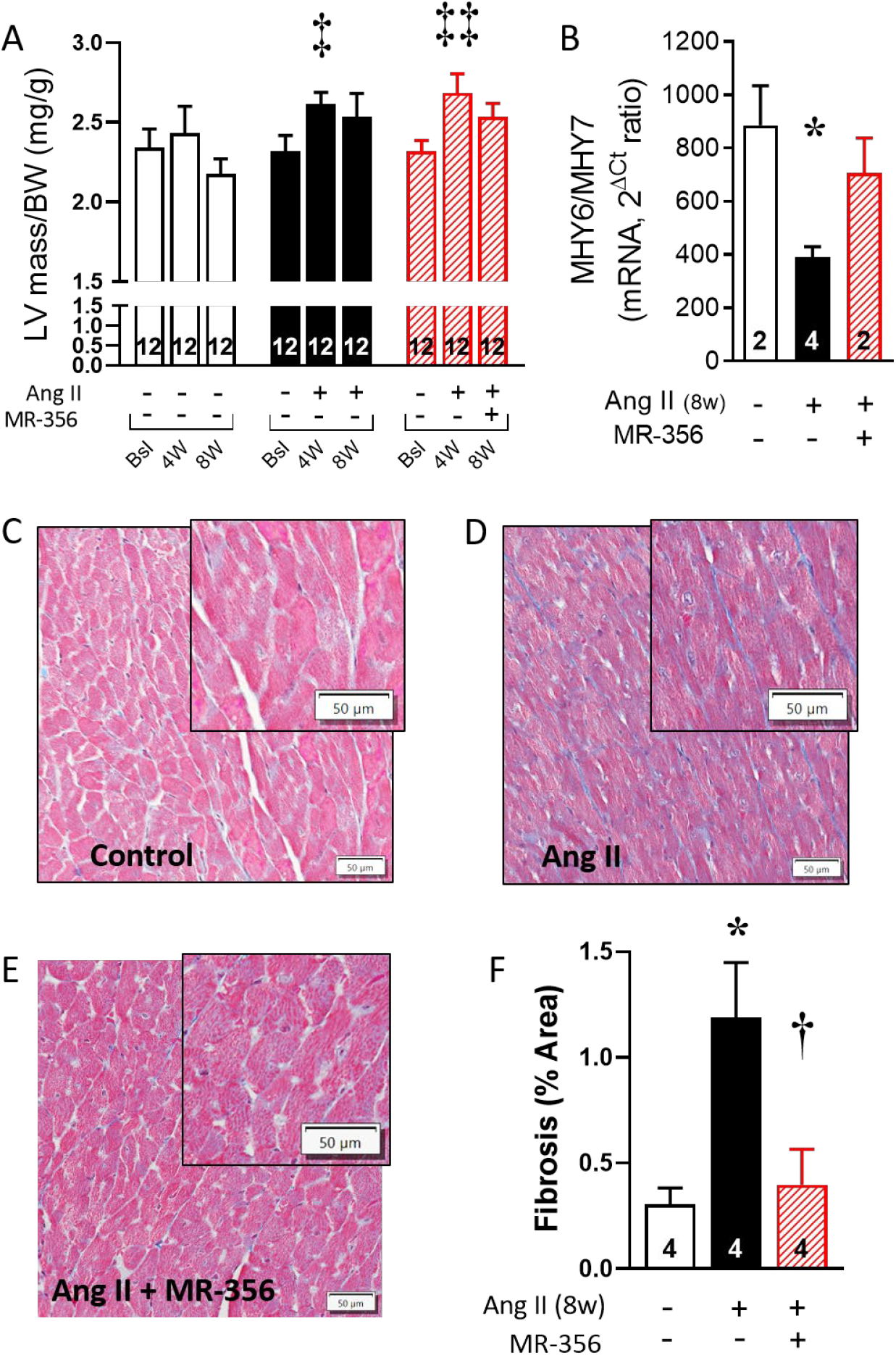
Myocardial remodeling is attenuated by treatment with GHRH-A. (A) LV mass assessed by echocardiography in control, Ang-II-infused (8-week) and Ang-II-infused (8-week)+MR-356 (4-week) mice assessed at baseline, and 4- and 8-weeks. (B) Gene expression ratio between αMHC (adult isoform) and βMHC (fetal isoform), as an indicator of the pathologic reprograming. (C) Representative Masson’s trichrome stained image of heart sections showing interstitial collagen deposition in control, (D) Ang-II (8-week) and (E) Ang-II (8-week)+MR-356 (4-week) mice. (F) Average percentage of fibrosis area. ^‡^*p*<0.05 and ^‡‡^*p*<0.01 vs. baseline, two-way ANOVA \**p*<0.05 vs. control, ^†^*p*<0.05 vs. Ang-II+MR-356-treated; one-way ANOVA.

In vivo hemodynamic assessments revealed that MR-356 restored EDPVR (Figure 8 A-D), as well as the relaxation time constant (τ) and the LVEDP (Figure 8E-F) to normal in HFpEF animals. The prolongation in IVRT during the first 4-week treatment with Ang-II in both groups, showed a continued increase in the Ang-II+placebo group but a substantial reversion in the MR-356 treated group (Figure 8G). The beneficial effects on diastolic function were also present in isolated cardiomyocytes, since MR-356 restored the resting sarcomere length (in diastole, Figure 8H, I). Moreover, sarcomere relaxation rate was accelerated by MR-356 treatment in comparison with the 8-week Ang-II group (Figure 8J). Treatment with MR-356 also reverted the delay in decay of Ca^2+^, consistent with the increased transcription (mRNA) of the SERCA2a gene (Supplemental Figure 4).

**Figure 8:**
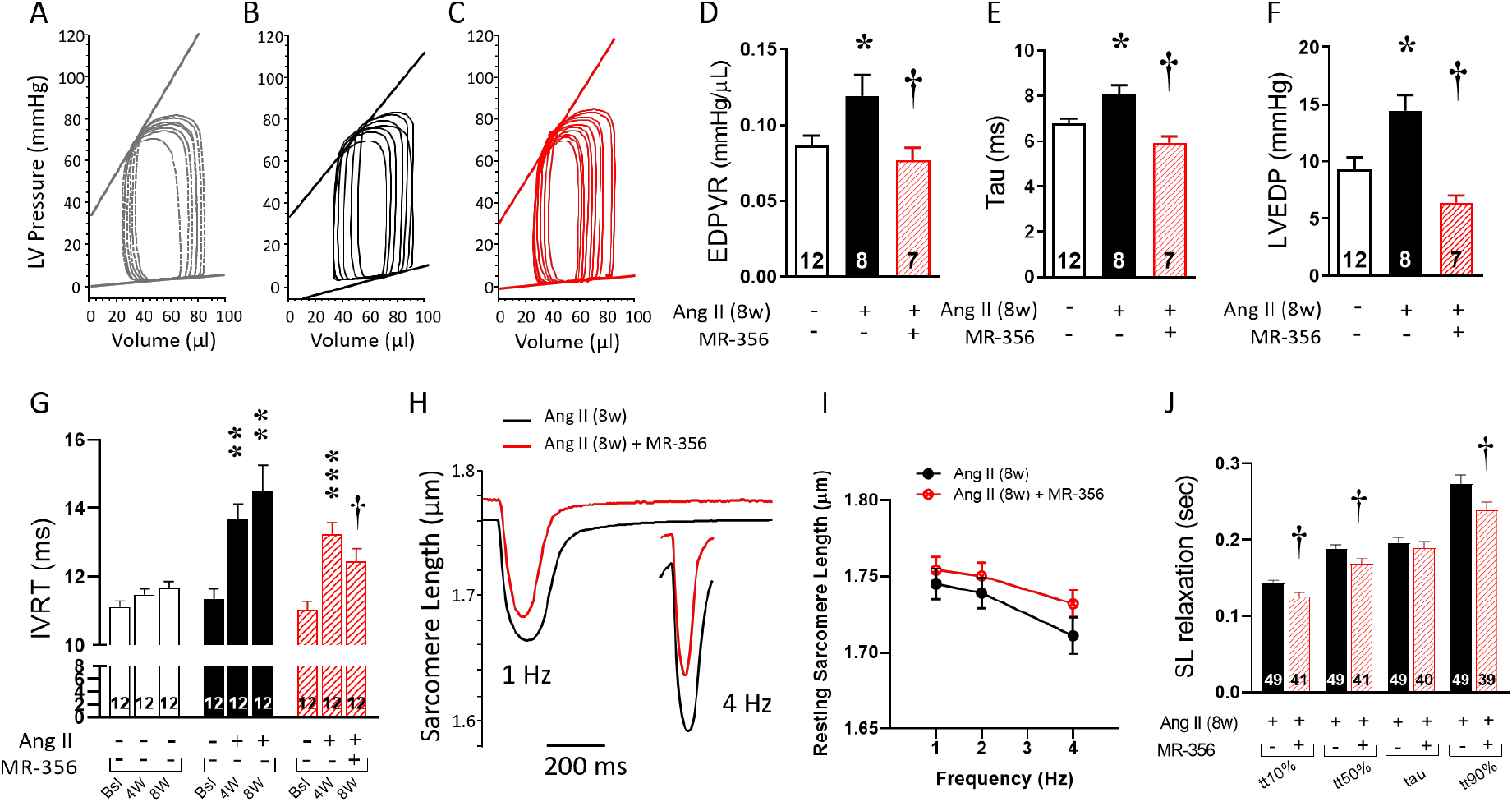
Effect of MR-356 on the chronic HFpEF phenotype. (A) Representative pressure-volume (P-V) loops showing the linear regression for ESPVR and EDPVR in control, (B) 8-week Ang-II-treated, or (C) Ang-II (8-weesk)+MR-356-(4-weeks)-treated mice. (D) Averaged slopes from the EDPVR linear fitting. (E) Hemodynamic assessment of the relaxation time constant (tau) at the 8-week time point. (F) Averaged LVEDP. (G) IVRT as measured by transthoracic echocardiography at baseline, and 4- and 8-weeks. (H) Representative SL traces comparing Ang-II (8-weeks)-treated and Ang-II (8-weeks)+MR-356 (4-weeks) cardiomyocytes at 1- and 4-Hz pacing rates. (I) Resting sarcomere length of Ang-II (8-weeks)-treated compared with Ang-II (8-weeks)+MR-356-treated (4-weeks) murine cardiomyocytes paced at 1-, 2- and 4Hz. (J) Averaged times to different degrees of SL relaxation: tt10%, tt50%, tau (τ) and tt90% in 8-week-treated cardiomyocytes paced at 1 Hz. ^†^*p*<0.05 vs. Ang-II (8-weeks)+MR-356-treated (4-weeks); Student’s t-test. ^‡^*p*<0.05 within group comparison, two-way ANOVA.

## Discussion

The incidence of the HFpEF syndrome is rising and it is unresponsive to conventional heart failure treatments. Given the rising clinical burden, it is imperative that effective treatments are found. Here we show that a novel class of peptide compounds, GHRH-As, previously shown to ameliorate pathological myocardial remodeling, also effectively restores diastolic dysfunction, a hallmark of HFpEF, to normal in this mouse model. GHRH-As, acts broadly through mechanisms involving fibrosis as well as myocyte calcium cycling and myofilament calcium response.

### In vivo effect of GHRH-A treatment

Chronic administration of Ang-II to mice reproduces structural and functional myocardial changes associated with HFpEF. In contrast to Regan el al.(27), our mice demonstrated hypertension, (aortic blood pressure was increased in Ang-II treated mice; Table 1). A majority of patients with HFpEF exhibit hypertension, which might represent a risk factor(34). These animals showed preserved EF and other contractility indices (although specifically ESPVR is reduced) but depressed overall function, as observed with GLS(33), associated with elevated LVEDP, myocardial fibrosis and impaired diastolic function, as measured by IVRT, EDPVR and the relaxation constant tau. Within this pathological context, HFpEF animals exhibited myocardial hypertrophy that was virtually redirected into a physiological hypertrophy by MR-356. Overall, this treatment improved myocardial performance associated with reduced indices of fibrosis, preserved cardiomyocyte width and diastolic RWT. However, MR-356 failed to prevent the increase in ventricle weight or reversion of the augmented LV mass (in the 8-week mice). The persistent myocardial hypertrophy after MR-356 treatment might be due to either effect, MR-356 directly activating GHRHRs on the myocardium, the action of GH (or IGF-1) released after MR-356 injections, or both (see below). Accordingly, our results show that the above-mentioned phenotype consistent with HFpEF syndrome, was both prevented and reversed by the treatment with the GHRH-A.

### GHRH-A targets myofilaments and calcium handling proteins

Isolated cardiomyocytes from HFpEF mice exhibited lusitropic impairment as well as alterations in Ca^2+^ handling, consistent with deficient myocardial relaxation and diastolic dysfunction. Our data strongly suggest that deficient myofilament relaxation in HFpEF cardiomyocytes is associated with higher affinity of myofilaments for Ca^2+^, favoring binding of cytosolic Ca^2+^ to TnC, thereby delaying cessation of the interaction between thin and thick filaments. This idea is supported by the finding of myofilament sensitization in HFpEF cardiomyocytes (leftward shift in the relaxation phase of SL-Ca^2+^ loops) associated with reduced phosphorylation of cTnI at serine 23/24 and cMyBPC at serine 282, which play important roles in regulating responsiveness of myofilaments to Ca^2+^(35,36). Ca^2+^ bound to myofilaments represents ~50% of the Ca^2+^ released from the SR in a typical heart beat(37). Therefore, increased affinity of myofilaments for Ca^2+^ has a substantial impact on cytosolic [Ca^2+^], being able to buffer large amounts of Ca^2+^, including the Ca^2+^ leaked from the SR in HFpEF cardiomyocytes, and therefore reducing the detectable levels. Moreover, the higher affinity for Ca^2+^ would make detachment from the troponin complex slower, impacting the cytosolic [Ca^2+^] decay rate(38,39). Phospholamban (PLB) exerts an inhibitory effect on SERCA2 by protein-protein interaction and PLB phosphorylation releases that repression on SERCA2. Increased PLB phosphorylation in HEpEF cardiomyocytes does not appear sufficient to compensate against both higher affinity for Ca^2+^ of myofilaments and diminished SERCA2 expression. Enhanced phosphorylation induced by MR-356 on PLB, together with its desensitizing effect on myofilaments, more efficiently counteracted and prevented slowing of Ca^2+^ decay rate (Figure 5B for the 4-week and Supplemental Figure 4A in the 8-week HFpEF models).

Myofilament responsiveness to Ca^2+^ was greatly decreased in cardiomyocytes from Ang-II+MR-356-treated mice. This lower affinity of myofilaments for Ca^2+^ might explain the molecular basis for faster relaxation, and may be why treatment with MR-356 preserved diastolic [Ca^2+^] in Ang-II-treated myocytes; i.e., preventing increased Ca^2+^ buffering capacity of myofilaments, since this molecular mechanism facilitates Ca^2+^ dissociation from cardiac myofilaments in response to increasing pacing(40). Phosphorylation of cTnI and cMyBPC accelerates relaxation either under β-adrenergic stimulation or increasing pacing frequency and may prevent diastolic dysfunction(41–44).

In addition to the underlying effect of elevated SRCa^2+^ leak on the depressed inotropic response of HFpEF cardiomyocytes, there may also be an inability to properly cycle cross-bridges. cMyBPC directly promotes force generation by enhancing cross-bridge binding and cooperative recruitment(45) and acts as a brake on cross-bridge kinetics by slowing down actin-myosin detachment(46), an effect removed by phosphorylation of cMyBPC (45,47,48). MR-356 treatment prevented depressed phosphorylation of cMyBPC at serine 282 in HFpEF cardiomyocytes and might account, in part, for the preserved contractile reserve in HFpEF cardiomyocytes treated with GHRH-A.

### Synthetic GHRH-As in animal models of cardiac diseases

MR-356 combines high GHRH activity with significant reduction of infarct size and prevention of ventricular remodeling prevention compared with other GHRH-As with high activation of GHRHR, but little or no myocardial repair capacity(17), possibly by stimulating reparative pathways including adenylate cyclase/PKA signaling as observed with GHRH(18). MR-356 also promotes survival of cardiac stem cell and induces proliferation in vitro, possibly mediated by ERK1/2 and PI3K/AKT pathways(49). A recent study using a model of pressure overload(22), which exhibits characteristics of HFrEF, shows that the GHRH-A MR-409 (another potent GHRH-A) attenuates hypertrophy and improves myocardial function. These beneficial effects are consistent with previous studies(19–21). JI-38, a GHRH-A comparable to MR-356 in terms of prevention of ventricular remodeling, promotes the up-regulation of the transcription factor GATA-4, mitosis and proliferation within the myocardium(19), which were not observed in rats treated with a recombinant GH, demonstrating that the direct actions of JI-38 on the myocardium are GH/IGF-1-independent. Despite the effectiveness of JI-38 in inducing GH secretion, circulating levels of GH were not different from control untreated animals. Thus, the transient increase of GH following MR-356 injection would likely not sustain long-term GH concentrations or maintain pathological levels. Our models of HFpEF exhibited ventricular hypertrophy that was not prevented or reversed by treatment with MR-356 (in contrast with Gesmundo’s publication(22)). This result might be due to a growth either in size or number of cardiomyocytes, or both. However, our data show that the cardiomyocyte size did not change. Moreover, MR-356 treatment induced an evident anti-fibrosis effect in our HFpEF mice. In contrast, among the important trophic role of GH/IGF-1 on the cardiovascular system(50), GH and IGF-1 promote fibrosis(51). Therefore, the observed effect would be more likely associated with a direct effect of the GHRH-A rather than with the GH/IGF-1 axis. This idea is supported by our previous findings of an GH/IGF-1-independent action of GHRH-As on the myocardium(19).

Our work focusses on a highly relevant clinical problem with major unmet needs, the HFpEF syndrome and the results presented here show for first time that GHRH-As can prevent/reverse the HFpEF phenotype, particularly diastolic dysfunction. Thus, based in the proven properties for cardiac repair of GHRH-As and the robust data obtained in the present study, we consider that GHRH-As are potentially useful for the management of HFpEF.

### Conclusion

In summary, we showed that activation of GHRHR with a synthetic GHRH-A prevents the appearance of HFpEF and reverses the HFpEF phenotype once established in mice, including dysfunctional diastolic relaxation in myocytes and intact animals and pathological myocardial remodeling. Increased SR Ca^2+^ leak associated with higher affinity of myofilaments for Ca^2+^ and impaired cross-bridge cycling, underlie the dysfunctional contractile response and lusitropy in these HFpEF cardiomyocytes, and may in part account for the alterations in Ca^2+^ kinetics. Myofibrillar proteins appear to be a target for treatment with GHRH agonists. Taken together these findings indicate that synthetic analogs of GHRH represent novel therapeutic compounds for treating patients with the HFpEF syndrome.

## Perspectives

### Clinical Competencies

HFpEF is arguably the major relevant clinical problem among the unmet medical needs currently affecting adults, and constitutes a leading cause of death and disability worldwide. Clinical trials have repeatedly failed to translate into HFpEF treatments, supporting the need for new approaches. Our study supports the potential use of synthetic GHRH-agonists that activate myocardial GHRH-receptor signaling pathways, as a novel therapeutic approach for HFpEF.

### Translational outlook

HFpEF is a clinical syndrome comprised of several pathologic conditions not limited to the heart. There are few animal models of HFpEF and none exhibits all aspects of this syndrome. The murine Ang-II-induced HFpEF model exhibits myocardial hypertrophy, fibrosis, aortic hypertension and diastolic dysfunction, with normal LV ejection fraction. Our study demonstrates a robust improvement in the relaxation indexes both in vitro and in vivo, including a clear antifibrotic effect. Clinical studies are needed to validate the effectiveness of GHRH-agonist treatment in humans.

## Visual abstract

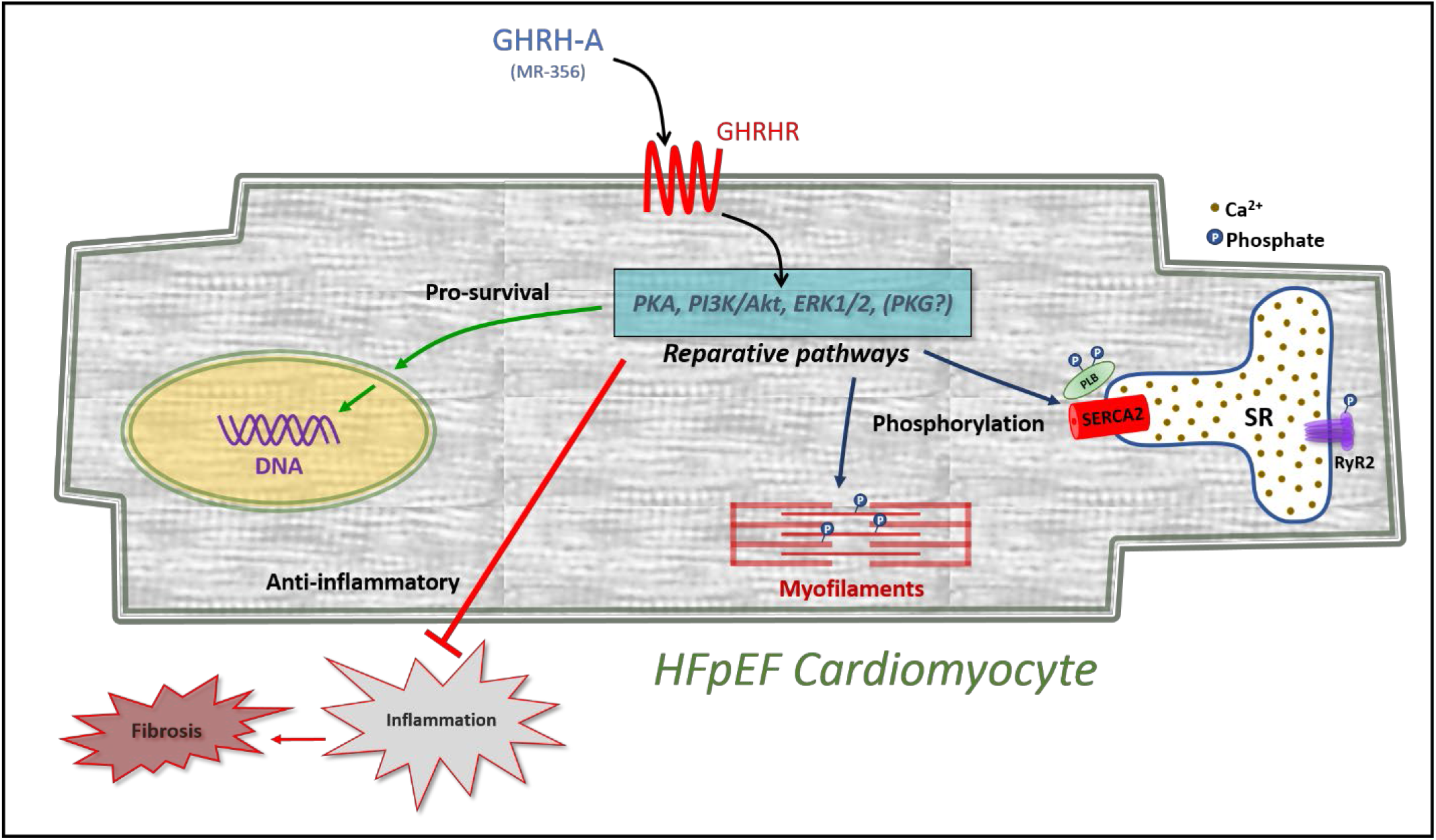
GHRH agonist as treatment for HFpEF: Schematic representation of the effects of a synthetic growth hormone-releasing hormone agonist (GHRH-A), MR-356, on a cardiomyocyte affected with heart failure with preserved ejection fraction (HFpEF). Activation of GHRH receptors (GHRHRs) triggers signaling pathways with reparative capacity. These pathways include phosphorylation of specific residues within myofibrilar proteins (e.g. cardiac TnI or cardiac MyBPC) and proteins of the calcium handling machinery, as well as activation of paracrine anti-inflammatory/anti-fibrotic signaling mechanisms.

## Supporting information

Expanded Methods and Supplemental Figures and Table

## Abbreviations

ABP: arterial blood pressure
Ang II: angiotensin II
BDM: 2,3-butanedione monoxime
BW: body weight
cMyBPC: cardiac myosin binding protein C
cTNI: cardiac troponin I
DMSO: dimethyl sulfoxide
dP/dt_max_: maximal contraction rate
dP/dt_min_: maximal relaxation rate
E: mitral peak velocity of early filling
Ea: arterial elastance
EDPVR: end-diastolic pressure-volume relationship
Ees: ventricular elastance
EF: ejection fraction
ESPVR: end-systolic pressure-volume relationship
E’: early diastolic mitral annular velocity
GAPDH: glyceraldehyde 3-phosphate dehydrogenase
GHRH: growth hormone-releasing hormone
GHRH-A: growth hormone-releasing hormone agonist
GHRHR: growth hormone-releasing hormone receptor
GLS: global longitudinal strain
HFpEF: heart failure with preserved ejection fraction
HFrEF: heart failure with reduced ejection fraction
HR: heart rate
Hsp90: heat shock protein 90
HW: heart weight
IVRT: isovolumic relaxation time
L-TCC: L-type Ca^2+^ channel
LV: left ventricle
LVEDD: left ventricle end-diastolic diameter
LVEDP: left ventricle end-diastolic pressure
LVEDV: left ventricle end-diastolic volume
LVESD: left ventricle end-systolic diameter
LVESV: left ventricle end-systolic volume
MyBPC: myosin-binding protein C
MHY: myosin heavy chain
NCX: Na^+^/Ca^2+^ exchanger
PLB: phospholamban
PRSW: preload recruitable stroke work
RWTd: relative wall thickness in diastole
SERCA2a: sarcoplasmic reticulum calcium ATPase 2a
SL: sarcomere length
SLS: sarcomere length shortening
SR: sarcoplasmic reticulum
Tau: relaxation constant
TGF-β: Transforming growth factor-β
cTnI: cardiac troponin I

## Acknowledgments

Sources of funding: This study was supported by NIH, grants 1R01 HL13735 and 1R01 HL107110 to JMH. JMH is also supported by NIH grants 5UM1 HL113460, 1R01 HL134558, 5R01 CA136387 and The Starr and Sofer Family Foundations. São Paulo Research Foundation (FAPESP) grant# 2016/01044-0 (to AGS).

