## Supplementary material for "Synthetic Growth Hormone-Releasing Hormone Agonist as Novel Treatment for Heart Failure with Preserved Ejection Fraction": Expanded Methods and Supplemental Figures and Table

**Supplemental Material**

**Expanded Methods**

*Experimental design and animal model*

We used age matched (2-month old) male CD1 mice (n=63, from Envigo). Mice were implanted with a mini-osmotic pump (Alzet) to deliver Ang-II (0.2 mg/kg/day; Sigma-Aldrich Co., Saint Louis, MO) for 4 weeks. Mice received daily injections of GHRH-Agonist (GHRH-A [MR-356]: 200 μg/kg, n=23) or vehicle (DMSO+propylene-glycol, n=23). Unmanipulated (no mini-osmotic pump) age-matched CD1 mice (n=17) were used as normal controls.

A separate group of male CD1 mice (n=24, 2-month old) was implanted with the Alzet mini-pumps containing Ang-II. After all mini-pumps content were delivered (4 weeks), the animals were subjected to replacement of mini-pumps for an additional 4-week Ang-II infusion. Upon the second set of mini-pump implantation (day 29), the mice were randomized. Half of them were daily injected with MR-356 and the remaining 12 mice received vehicle injections.

*Materials*

The GHRH-A, MR-356 ([N-Me-$Tyr^1$, $Gln^8$, $Orn^{12}$, $Abu^{15}$, $Orn^{21}$, $Nle^{27}$, $Asp^{28}$, $Agm^{29}$] hGH-RH(1-29)), was synthetized by solid phase methods and purified by HPLC in our laboratories as described previously(1).

*Echocardiography*

Transthoracic echocardiography was performed using a Vevo 2100 ultrasound system equipped with a 30 MHz transducer (MS400) (Visual Sonics Inc., Toronto, Ontario, Canada) as previously described with minor modifications(2). Briefly, mice were anesthetized with isoflurane (3%, induction chamber) and maintained with 0.5-2% using a facemask with a scavenging system. Each imaging sessions took around ~10-15 minutes. The echocardiograms were used to image non-invasively across the mouse's skin, which were cleared of fur at the site using a depilatory cream. Ultrasound gel were used as a coupling fluid between the transducers and the skin. Following echo, animals were allowed to regain consciousness. Animals were directly monitored during this period to insure full recovery. Mice were screened at baseline and 4 weeks following mini-osmotic pump placement, as well as at 8 weeks (final time point). A parasternal long-axis and short-axis views were acquired during the exam for cardiac morphology and function. In addition, Doppler mitral blood flow were obtained to assess diastolic function. All echocardiographic measurements were performed offline over 3 to 5 heart beats.

*Speckle Tracking Echocardiography (STE)*

Strain analysis was conducted using speckle tracking software, Vevostrain TM analysis (Visual Sonics). Global longitudinal strain was calculated using B-mode cine images in the LV parasternal long-axis view acquired at a high frame rate(3).

*Hemodynamic Measurements*

Hemodynamic studies were performed using a micro-tipped pressure-volume catheter (SPR-839; Millar Instruments) as previously described with minor modifications(4,5). Briefly, mice were anesthetized with isoflurane (3% induction chamber). Body temperature was controlled (~37°C) during the whole procedure. When anesthetic action started, the animal was transferred to the surgical bench and anesthesia maintained with ~2% Isoflurane. Hair remover (e.g., Nair) was spread on the animal's chest to remove the fur and iodo-alcohol solution applied to establish an aseptic field. A small incision was made in the skin over the neck to permit the endotracheal intubation (through the mouth). The left internal jugular vein was exposed and cannulated with a 30-gauge needle for the administration of drugs or fluid support. The skin over the site of median ventral neck was cut and the carotid artery exposed. After occlusion of the distal portion of the carotid artery, a small cut in artery was done to permit the catheter being inserted and advance retrograde into the LV. The catheter is equipped with a pressure transducer and a sensor for measuring chamber conductance to estimate the ventricular volume. The recorded conductance is calibrated offline by using diastolic and systolic ventricular volumes obtained by echocardiography. P-V loops were recorded during steady-state and temporary inferior vena cava (ivc) occlusion. Occlusions of ivc were performed by pushing the belly with a cotton tip or by direct occlusion using a small forceps after opening the chest (~4-5th intercostal space). The ventilator was stopped momentarily (~10 s) to avoid breathing interference during measurements. At the end of the experiments, the animals (under deep anesthesia as above) were humanely euthanized and the hearts were harvested for molecular or in-vitro studies. All analyses were performed using LabChart Pro version 8.1.5 software (ADInstruments). The software was configured to plotting P-V loops; end-systolic pressure versus end-systolic volume values were fit using a linear regression (least squares), and the end-diastolic pressure versus end-diastolic volume values were fit using either a linear regression, based on the following model:

Linear equation $\boldsymbol{P} = \boldsymbol{A} + \boldsymbol{B} * \boldsymbol{V}$,

where $P$ is pressure (mmHg) and $V$ is volume (μL). $A$ is the ordinate constant (mmHg). $B$ is the slope that represents the "linear stiffness coefficient" (mmHg/μL). The reported data expressed as EDPVR (mmHg/μL) correspond to $B$, the "stiffness coefficient".

*Cardiomyocyte Isolation*

Cardiac myocytes were isolated and prepared from hearts of mice as described previously(6). Briefly, hearts were harvested and retrograde perfused through the aorta (at constant flow 2 mL/minutes) in a modified Langerdorf system with an isolation solution containing (in *mmol/L*): *120* NaCl, *5.4* KCl, *1.4* $MgSO_4$, *1.2* NaH2PO4, *20* $NaHCO_3$, *10* 2,3-butadiene monoxime (BDM, Sigma-Aldrich Co.. Saint Lois, MO), *5* taurine (Sigma-Aldrich Co. Saint Lois, MO), *5.6* glucose (bubbled with 5% $CO_2$ - 95% $O_2$ for at least 15 minutes). After 5-10 minutes blood flushing at 37°C, perfusing solution was switched to a one with collagenase type 2 (Worthington Biochemical Corporation, Lakewood, NJ) ~315 U/mL and protease type XIV (Sigma-Aldrich Co., Saint Louis, MO) 5.2 U/mL. Then, digested hearts were chopped and myocytes were released by gentle mechanical disruption, filtered through a 500 nm cell strainer and gently pulled down by centrifugation at 800 rpm for 1 minute. Cardiomyocytes were resuspended in isolation solution containing BDM and 0.5 % bovine serum albumin (BSA, Sigma-Aldrich Co. Saint Lois, MO). A small portion of this cell suspension was kept for cell length and width measurement, and the rest was subjected to a sequential extracellular $Ca^{2+}$ restoration on a Tyrode's buffer containing (in *mmol/L*): *144* NaCl, *1* $MgCl_2$, *10* HEPES, *5.6* glucose, *5* KCl, *1.2* $NaH_2PO_4$ (adjusted to a pH 7.4 with NaOH). Finally, cardiomyocytes were resuspended in Tyrode's buffer containing 1.8 mmol/L $CaCl_2$ at room temperature.

*Intracellular Calcium and Sarcomere Length Measurement*

Intracellular $Ca^{2+}$ was measured using the $Ca^{2+}$-sensitive dye Fura-2 and a dual-excitation (340/380 nm) spectrofluorometer (IonOptix LLC, Milton, MA, USA). First, cardiomyocytes were incubated with 2.5 μmol/L Fura-2 for 15 minutes at room temperature and then washed with fresh regular Tyrode's solution for at least 10 minutes. Then, cardiomyocytes were placed in a perfusion chamber adapted to the stage of an inverted Nikon eclipse TE2000-U fluorescence microscope. Cells were superfused with a Tyrode's buffer at 37°C and Fura-2 fluorescence was acquired at an emission wave length of 515±10 nm. The cardiomyocytes were electric field-paced (20V) at different frequencies from 0.5 to 4 Hz (depending on the protocol). The calibration was performed in cardiomyocytes "ex vivo" superfusing a free $Ca^{2+}$ and then a $Ca^{2+}$ saturating (5 mmol/L) solutions, both containing 10 µmol/L ionomycin (Sigma, St. Louis, MO) until reaching a minimal ($R_{min}$) or a maximal ($R_{max}$) ratio values, respectively. $[Ca^{2+}]_i$ was calculated as described previously(6) using the following equation:

$$[\boldsymbol{Ca^{2+}}]_{\boldsymbol{i}} = K_d \times \frac{S_{f2}}{S_{b2}} \times \frac{(\mathrm{R} - R_{\mathrm{min}})}{(R_{max} - \mathrm{R})}$$

$K_d$ (dissociation constant) in adult myocytes was taken as 224 nmol/L. The scaling factors $S_{f2}$ and $S_{b2}$ were extracted from calibration as described by Grynkiewicz et al(7). $\Delta[Ca^{+2}]_i$ amplitude was considered as: peak $[Ca^{+2}]_i$ – resting $[Ca^{+2}]_i$.

Simultaneously, sarcomere length (SL) was recorded with an IonOptix iCCD camera. Change in average SL was determined by Fast Fourier Transform analysis of the Z-line density trace to the frequency domain, and SL shortening was calculated as follows:

$$\boldsymbol{SL\ Shortening}\ (\%) = 100 \times \frac{(resting\ SL - peak\ SL)}{resting\ SL}$$

*Hysteresis Sarcomere shortening-Calcium loops analysis*

The hysteresis loops were obtained by plotting sarcomere length versus $Ca^{2+}$ concentration. Two of the three main segments of such loops were analyzed for this work. The segment transitioning between the maximum $[Ca^{2+}]$ (point B on Supplemental image 1 below) and the maximum shortening (peak contraction, point C on Supplemental image 1) correspond to the "force-activation phase". The "relaxation phase" transitions between this maximum shortening (point C) and the point where the minimum shortening and minimum $[Ca^{2+}]$ merge (point A on Supplemental image 1). The analysis of the relaxation phase (C-A) was assessed by a non-linear (sigmoidal) fit, while the force-activation phase (B-C) was subjected to a linear regression(8,9).

Both analyses were performed by GraphPad Prism version 8.3.0 (GraphPad Prism Software Corporation San Diego, CA USA).

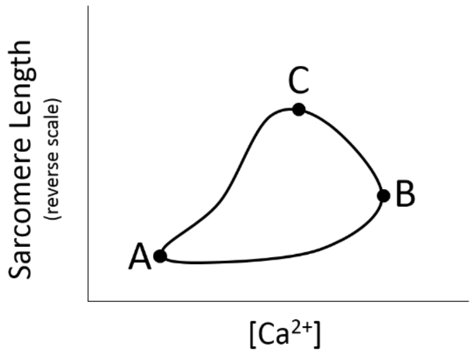


Schematic representation of a typical SL-[$Ca^{2+}$] loop depicting the three most relevant spots: A, resting; B, maximum [$Ca^{2+}$]; C, maximum sarcomere shortening.

*SR$Ca^{2+}$ Leak and SR $Ca^{2+}$ Load Measuring*

SR$Ca^{2+}$ leakage was assessed using tetracaine by a modified protocol from the one described by Shannon *et al*(10). Briefly, after stop pacing the cardiomyocytes, the superfusing buffer was immediately switched for a 0$Na^+$/0$Ca^{2+}$ Tyrode's buffer containing (*in mmol/L*): *144* LiCl, *1* $MgCl_2$, *10* HEPES, *5.6* glucose, *5* KCl, *10* EGTA (pH adjusted up to 7.4 with LiOH) during approximately 30 seconds to prevent $Ca^{2+}$ fluxes through the sarcolemma. Thus, the intracellular $Ca^{2+}$ is maintained approximately unaltered and the cytosolic [$Ca^{2+}$] reaches a steady state due to a balance between the SR$Ca^{2+}$ fluxes ($Ca^{2+}$ uptake and $Ca^{2+}$ leak). Then a 0$Na^+$/0$Ca^{2+}$ Tyrode's solution with 1 mM tetracaine (Sigma-Aldrich) was applied for at least 60 seconds. Under this condition, the SR$Ca^{2+}$ leak is blocked and then allowing the $Ca^{2+}$ uptake to reduce cytosolic [$Ca^{2+}$] until reaching a minimum in the Fura-2 ratio. Following, the superfusing solution was switched back to tetraine-free 0$Na^+$/0$Ca^{2+}$ Tyrode's buffer and SR$Ca^{2+}$ content was assessed as described by Bassani *et al*(11) by a 20 mmol/L caffeine challenge. SR$Ca^{2+}$ contents were calculated considering that SR represents 3.5% and cytosol 65% of the myocyte volume as previously described(10). The following equation from Shannon et al. was used:

$$\boldsymbol{SR}[\boldsymbol{Ca^{2+}}] = [Ca^{2+}]_{caff} + \frac{\beta_{max-SR} \times [Ca^{2+}]_{caff}}{[Ca^{2+}]_{caff} + K_{d-SR}}$$

SR[$Ca^{2+}$] is the $SRCa^{2+}$ content, $[Ca^{2+}]_{caff}$ is the SR $Ca^{2+}$ released by caffeine, $\beta_{max-SR}$ and $K_{d-SR}$ are the usual Michaelis parameters for SR $Ca^{2+}$ binding.

The observed decrease in the Fura-2 ratio in presence of tetracaine compared to the ratio before applying tetracaine was considered the $Ca^{2+}$ leak for a particular myocyte (Figure 3A; expressed in μmol/L). $SRCa^{2+}$ leak-SR[$Ca^{2+}$] pairs were grouped by similar $SRCa^{2+}$ load (SR[$Ca^{2+}$]), plotted and fitted by an exponential growth function using the Graph Pad Prism software (version 8.4.3) (Figure 4B). Additionally, $SRCa^{2+}$ leak values corresponding to SR[$Ca^{2+}$] averaging 100 μmol/L were plotted in a bar graph for a clearer appreciation of the data (Figure 3C). Measurements were carried out at 37°C.

*Western Blot Immunoanalysis*

*Isolated Cardiomyocytes:* Frozen cardiomyocytes were lysed in ice-cold RIPA buffer with proteinase inhibitors (cOmplete protease inhibitor cocktail; Roche), sonicated and then centrifuged at 12500 RPM for 15 minutes at 4°C. Supernatants were collected and sampled for protein concentration by BCA assay (Thermo Scientific). Samples (40μg of protein from myocyte lysates) were electrophoresed using a NuPAGE 10% or 15% Bis-Tris gels (Invitrogen) and transferred to PVDF or nitrocellulose membranes (Bio-Rad Laboratories) as convenient. Immunoblot analysis was performed with antibodies against cardiac troponin I (cTnI, Abcam), phospho-serine 23/24 cTnI (Cell Signaling Technology, #4004S), the cardiac sarcoplasmic reticulum calcium ATPase, SERCA2a (Santa Cruz Biotechnology, Inc., # sc-8095), phospholamban (PLB; Pierce, Thermo Scientific™, # MA3-922), phospho-serine 16 PLB (Badrilla, Leeds, UK, # A010-12), phospho-threonine 17 PLB (Badrilla, Leeds, UK, # A010-13), MyBPC3 (Santa Cruz Biotechnology, Inc., #sc-137237), phospho-serine 282 cardiac MyBPC (Enzo Life Sciences, # ALX-215-057-R050) and Hsp90 (Cell Signaling Technology, # 4874S)(overnight incubation at 4°C). As fibrosis/inflammation biomarker, a polyclonal anti-TGF-β (Cell Signaling, #3711) antibody was used. The following secondary antibodies horse radish peroxidase (HRP)-conjugated were used: anti-rabbit IgG (Cell Signaling Technology, #7074), anti-mouse IgG (Cell Signaling Technology, #7076) and anti-goat IgG (Santa Cruz, sc-2350), and incubated for 1 hour at room temperature. Antibodies' dilutions were adjusted within the range recommended by the vendor. The signal was visualized using the SuperSignal™ West Femto Maximum Sensitivity Substrate (Thermo Scientific™, # 34094). Phosphorylated PLB was expressed as a ratio of p-Ser16/PLB or p-Thr17/PLB, phosphorylated cTnI was expressed as p-Ser23-24/cTnI or p-Ser43/cTnI, phosphorylated cMyBPC was expressed as p-Ser282/cMyBPC. Expression of proteins were compared with the level loading control, Hsp90 or β-actin (Santa Cruz Biotechnology, Inc. #sc-47778). Images were analyzed by the ImageJ software (developed by the National Institutes of Health).

*Hearts:* Frozen cardiac tissues were homogenized using a Pyrex glass-glass homogenizer on liquid nitrogen and urea buffer containing: 8 mol/L urea, 2 mol/L thiourea, 3% SDS, 0.03% bromophenol blue and 0.05 mol/L Tris, pH 6.8. Glycerol (50%) was added. The samples were then centrifuged at 12500 RPM for 5 minutes. Supernatants were collected and sampled for protein concentration by Bradford assay (Thermo Scientific). Samples (40μg of protein) were loaded on a 10% agarose gel, electrophoresed and transferred to PVDF or nitrocellulose membranes (Bio-Rad Laboratories) as convenient. Membranes were blocked with 1% Tween-TBS + Rockland blocking buffer MB-070 (1:1). Immunoblot analysis was performed using rabbit polyclonal antibodies against collagen I (Abcam, #34710) and collagen III (Abcam, #7778), and mouse monoclonal antibody for GAPDH (Cell Signaling, #97166S) as internal loading control (overnight incubation at 4°C). Detection was carried out by incubating with the fluorescent-dye conjugated antibodies anti-rabbit (IRDye 680RD, LiCor, # 925-68071) and anti-mouse IgG (IRDye 800CW, LiCor, # 925-32210) for 1 hour at room temperature. Images were acquired by an Odyssey infrared imaging system (Li-Cor Biosciences, Lincoln, NE) and then analyzed by the ImageJ software (NIH). As fibrosis biomarker, an anti-prolyl 4-hydroxylase α1 (Proteintech Group Inc., Rosemont, IL, #12658-1-AP) antibody was normalized by the loading control, Hsp90. Detection was performed by using the HRP-conjugated secondary antibody indicated above, and chemiluminescence signal was acquired and analyzed as described.

*Interstitial Fibrosis Quantification*

Hearts were harvested and fixed in 4% paraformaldehyde. Each heart was dehydrated with graded alcohols, embedded in paraffin and longitudinally sliced into 5-μm sections. The sections were subjected to Masson's trichrome staining for assessment of interstitial fibrillary collagen deposition. Images were acquired by an Olympus VS120 scanner (Olympus

Corporation, Life Science Solutions) and analyzed with the ImageJ software (by the National Institutes of Health) by an investigator blinded to the groups.

*Quantitative RT-PCR*

Total RNA was isolated from cardiomyocytes using RNAeasy mini kit (Qiagen, Hilden, Germany). RNA (1 µg) was reverse-transcribed according to instructions utilizing a cDNA Synthesis kit (Applied Biosystems, Foster City, CA). Using Taqman® Universal Master Mix (Applied Biosystems, Foster City, CA) in a Bio-Rad CFX96™ Real-Time PCR system (Bio-Rad Laboratories, Hercules, CA, USA), qPCR was performed. All samples were run in duplicates and normalized to 18S Taqman gene-expression as follows: ATP2a2: Mm01201431_m1; RYR2 Mm00465877_m1, TNNi1: Mm00502426_m1; TNNi3: Mm00437164_m1; MHY6: Mm00440359_m1; MHY7: Mm00600555_m1and 18S Mm03928990_g1. The change in mRNA expression was calculated compared to the change in normalized 18S, and values were expressed as 2ΔCt.

*Inclusion and Exclusion criteria*

All animals that completed the protocols and subjected to in vivo ultrasound evaluation were included in the study. However, missing data for some parameters is due to technical issues in either the acquisition or the measurements of the records. Pressure-volume (P-V) hemodynamic measurements were performed in a subset of animals (from the 4-week study) which tolerated the anesthesia for the length of the procedure (n=8 control, n=9 Ang II-treated and n=10 Ang II+MR-356-treated mice were subjected to hemodynamic assessment). Of these, 1 outlier [beyond 2x SD units) animal from the Ang-II+MR-356 group was excluded). In the hemodynamic assessment from the 8-week study (n=12 per group), data were excluded (n=4 Ang-II (8w) and n=5 Ang-II (8w)+MR-356) due to either outlier values (beyond 2x SD units) or inappropriate quality in P-V hemodynamic records. The criterion for exclusion of any sample of isolated cardiomyocytes was the expert evaluation about the quality of the whole batch of isolated cells. On the other hand, samples for western blot analysis were excluded only if the quantitative yielding was insufficient (independently of the quality of cells).

*Statistical Analysis*

Data are reported as mean ± standard error of the mean (SEM). Statistical significance between three groups was determined by one-way ANOVA, or two-way ANOVA followed by Tukey's or Bonferonni's post hoc tests, as appropriate. For comparisons of two groups, Student's t-test was used. Analyses were performed using GraphPad Prism version 8.4.3 (GraphPad Software Corporation, San Diego, CA USA). The null hypothesis was rejected at $p<0.05$.

*Study approval*

All protocols and experimental procedures were approved by the University of Miami Animal Care and Use Committee following the Guide for the Care and Use of Laboratory Animals (NIH Publication No. 85-234, revised 2011).

**Supplemental Figures:**

- Supplemental Figure 1: Cardiomyocytes length in full-relaxed conditions.

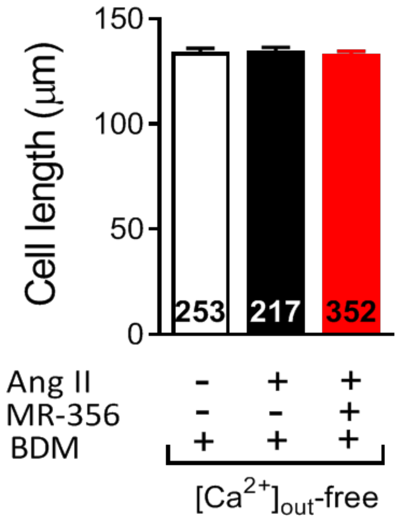


**Supplemental Figure 1: Cardiomyocytes length in full-relaxed conditions.** Quiescent cardiomyocytes length in $Ca^{2+}$-free and the presence of BDM buffer (Actual length of completely relaxed cardiomyocytes where myofilament interaction is uncoupled). Numbers on bottom of bars correspond to analyzed cardiomyocytes from N=4 control, N=5 Ang-II-treated and N=5 Ang-II+MR-356-tretaed mice.

- Supplemental Figure 2: Additional fibrosis/remodeling markers.

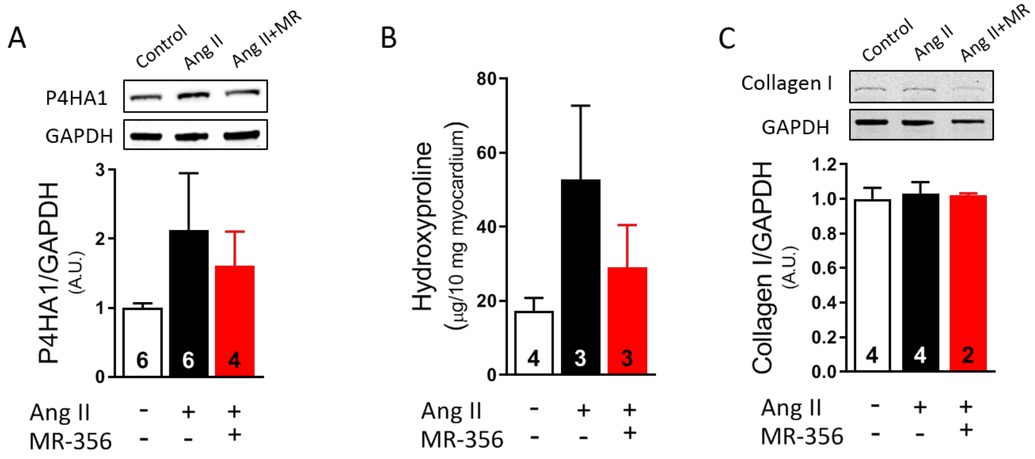


**Supplemental Figure 2: Additional fibrosis/remodeling markers**. (A) Prolyl 4-hydroxylase subunit α-1 (P4HA1) protein expression. P4HA1is a key enzyme for the hydroxylation of proline residues on collagen. (B) Hydroxyproline content. Hydroxyproline is a modified amino acid residue characteristic of the collagen primary structure. (C) Collagen I protein expression in heart tissue. Numbers on bottom of bars correspond to analyzed heart homogenates. Data were subjected to one-way ANOVA.

- Supplemental Figure 3: Hypercontracted cardiomyocytes and depressed calcium-frequency response in the HFpEF model.

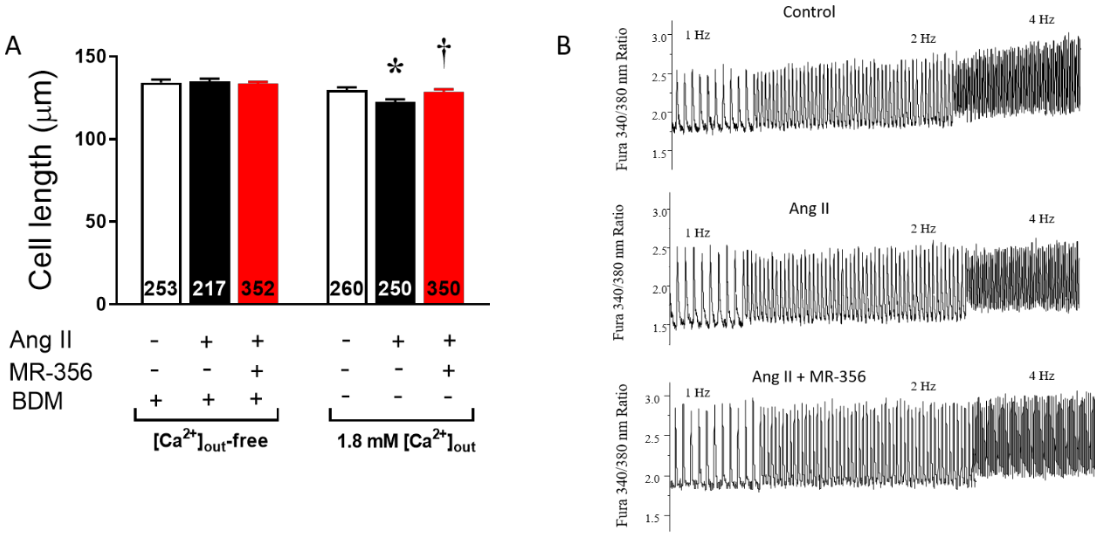


**Supplemental Figure 3: Hypercontracted cardiomyocytes and depressed calcium-frequency response in the HFpEF model.** (A) Cardiomyocyte length in $Ca^{2+}$-free and the presence of BDM buffer (left, where myofilament interaction is uncoupled), or in the presence of $Ca^{2+}$ and the absence of BDM (right, normal conditions where myofilaments are able to interact). Numbers on bottom of bars correspond to analyzed cardiomyocytes from N=4 control, N=5 Ang-II-treated and N=5 Ang-II+MR-356-tretaed mice. * $p<0.05$ vs. control and † $p<0.05$ vs. Ang-II-treated; one-way ANOVA. (B) Representative $Ca^{2+}$ transients in cardiomyocytes from untreated control (N=4), Ang-II (N=5) or Ang-II+MR-356-treated (N=5) mice, paced at 1, 2 and 4 Hz.

- Supplemental Figure 4: GHRH-A improves SR $Ca^{2+}$ re-uptake in cardiomyocytes from the 8-week Ang-II infusion mouse model.

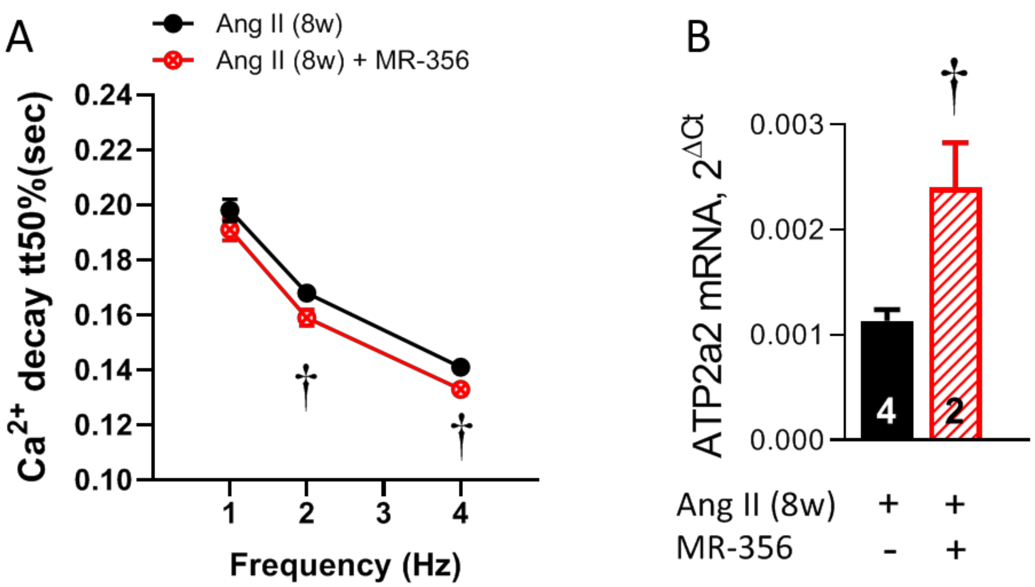


**Supplemental Figure 4: GHRH-A MR-356 improves SR $Ca^{2+}$ re-uptake in cardiomyocytes from the 8-week Ang-II infusion mouse model.** (A) Time to 50% $Ca^{2+}$ decay at 1, 2 and 4 Hz electric-field stimulation in cardiomyocytes from Ang-II (8w)-infused compared to Ang-II (8w)+MR-356-treated mice. (B) SERCA2a gene expression. Increase in SERCA2a expression induced by MR-356 in HFpEF cardiomyocytes consistently correlates with faster $Ca^{2+}$ decay. † $p$<0.05 vs. Ang-II (8w)-treated; two-way ANOVA or Student's t-test as correspond.

**Supplemental tables:**

- Supplemental Table 1: Biometric values of 4-week model.

| | Control (N=15) | Ang-II (N=9) | Ang-II+MR-356 (N=9) |
|---|---|---|---|
| **HW (mg)** | 182.1±4.8 | 205.6±9.8* | 210.1±8.9* |
| **BW (g)** | 44.0±0.9 | 41.9±1.3 | 42.6±0.8 |
| **TL (mm)** | 18.73±0.08 | 18.97±0.15 | 18.94±0.13 |
| **HW/BW (mg/g)** | 4.14±0.07 | 4.89±0.13* | 4.94±0.21* |
| **HW/TL (mg/mm)** | 9.7±0.2 | 10.8±0.5* | 11.1±0.5* |

**Supplemental Table 1:** All values represent mean±SEM. HW, heart weight; BW, body weight; TL, tibia length; * $p<0.05$ vs. control; † $p<0.05$ Ang-II+MR-356 vs. Ang-II, one-way ANOVA. N represents number of studied animals.
